# Improved transfection methods of primary cultured astrocytes for observation of cytoskeletal structures

**DOI:** 10.1101/2025.01.27.635031

**Authors:** Chieko Ikoma, Kodai Inoue, Kouta Kasai, Satoko Tsukuda, Akiko Tamura, Shihoko Nakata, Yuto Iwata, Takumi Tamagawa, Ayako Nakayama, Kazunori Takano, Eiji Shigetomi, Schuichi Koizumi, Hiroyuki Nakagawa, Asako G Terasaki

**Author notes:** These authors contributed equally to this work (Chieko Ikoma, Kodai Inoue).

## Abstract

Astrocytes are the predominant type of glia in the central nervous system and have long-branched stem processes and perisynaptic/peripheral astrocyte processes (PAPs) contacting neurons and other glial cells. However, a common astrocyte culture method generated undesired fibroblast-like cells; thus, the roles of cytoskeletal proteins in astrocytes have not been well studied. Previously, we reported a culture method of chicken astrocytes forming structures similar to stem processes and PAPs in vivo. In the current study, we improved transfection methods retaining astrocyte morphology at low cell density, suitable for observing protein behaviors. Our cultured astrocytes had various actin-containing substructures such as filopodia, lamellipodia, and microvilli in actively moving PAP-like structures. Moreover, plasma membrane-actin linking protein ezrin (a PAP marker in brain tissues) and lasp-2 (LIM and SH3 protein 2, highly expressed in cultured astrocytes) accumulated in the different actin-containing substructures. Additionally, lasp-2 and F-actin colocalized as small elliptical structures at the base of lamellipodia and filopodia of process tips, which may be cell-substrate adhesions. Our developed methods offer significant advantages for analyzing the regulation of astrocyte morphology and motility.

## INTRODUCTION

In the nervous system, neuronal function is maintained by various kinds of glial cells. Astrocytes are the predominant type of glia in the central nervous system (CNS). The term “astrocyte” is derived from stellate (aster, star-like) morphology with radial processes expressing intermediate filaments of glial fibrillary acidic protein (GFAP) observed in vivo. GFAP is a common astrocyte marker; in rodent brains, there are two main GFAP-positive astrocyte types: fibrous astrocytes organized along white matter tracts and protoplasmic astrocytes distributed relatively uniformly within cortical gray matter (reviewed by Ben Haim & Rowitch, 2016). Mature protoplasmic astrocytes comprise a cell body, several major branches from the cell body referred to as “stem processes”, and “perisynaptic / peripheral astrocyte processes” (PAPs) extending from the stem processes. Some dyes, such as Lucifer yellow, can visualize whole astrocytes in tissues with bushy processes (Moye et al., 2019). Each astrocyte covers a large nonoverlapping territory with neighboring astrocytes (typically 50-100 μm diameter in mice and 100-400 μm in humans) (Oberheim et al., 2009), and each territory is thought to be a functional compartment (Endo et al., 2022). PAPs are known to regulate neuronal excitability and synaptic transmission via formation of tripartite synapses (reviewed by Semyanov & Verkhratsky, 2021). Astrocyte-neuron interactions also contribute to neural development and differentiation (reviewed by Ben Haim & Rowitch, 2016). Astrocytes also enwrap layers of brain endothelial cells and form the blood-brain barrier with “endfeet”, which are specialized distal extensions of astrocytes that contact the vasculature. Recently, astrocytes have been revealed to form astrocyte-oligodendrocyte and astrocyte-astrocyte interactions (reviewed by Yu & Khakh, 2022).

Historically, astrocyte morphological heterogeneity within brain areas of mammalian tissues has been reported, and this may be related to astrocyte functional heterogeneity. During brain development, astrocytes arise from glial progenitors, disperse across cortical cell layers and columns, and increase region-specific morphological complexity (reviewed by Torres-Ceja & Olsen, 2022). Astrocytes can also become “reactive astrocytes” responding to CNS damage and disease, involving morphological changes including cell body hypertrophy and thickening of the major processes, along with actin and GFAP overexpression (reviewed by Lawrence et al., 2023). The morphology of astrocytes and neurons is highly relevant to brain function (reviewed by Wilson et al., 2023), and some animal disease models showed morphological abnormality of astrocytes (reviewed by Baldwin et al., 2024). For example, territory areas of astrocytes were smaller in model mice of Huntington’s disease, a severe neurodegenerative disease caused by a polyglutamine-encoding CAG expansion in the huntingtin (Htt) gene (Octeau et al., 2018). Also, in SAPAP3 knockout mice showing obsessive-compulsive disorder phenotypes, reduced territory sizes and disrupted actin cytoskeletal organization were observed (Soto et al., 2024).

In many cell types, adhesion, contraction, migration, and shape changes depend on actin cytoskeleton (reviewed by Svitkina, 2018). Various actin-binding proteins have been reported to control actin filament dynamics in the formation of actin-containing structures such as lamellipodia, filopodia, focal adhesions, and adherens junctions. Indeed, PAPs are observed as lamellate processes and filopodia enriched in actin microfilaments in brain tissues, and the actin-binding protein ezrin, connecting F-actin to the cell membrane, was found to be localized at the neuropil regions of rat hippocampus (considered to comprise PAPs etc.) (Derouiche & Frotscher, 2001). Additionally, ezrin was also observed across the entire surface of morphologically intact astrocytes immediately after dissociation from the murine cerebral cortex (Haseleu et al., 2013). However, the mechanisms by which actin-binding proteins regulate migration during development, branch emergence, PAP formation, enwrapping of synapses, and connections with other glial cells remain poorly understood.

One reason for this is that astrocytes in vivo have complex 3D structures as described above. Another reason is the culture conditions of primary astrocytes. A common method for culturing astrocytes in conventional Dulbecco’s Modified Eagle Medium (DMEM) supplemented with fetal bovine serum (FBS) generated undesired fibroblast-like cells without stem processes (Aumann et al., 2017). Despite morphological differences, cultured fibroblast-like astrocytes expressed GFAP and many astrocytic functions, such as metabolism, neurotransmission, and calcium signaling, were also observed in cultured cells (reviewed by Lange et al., 2012). Moreover, standard culture protocols using conventional medium recommend high cell density, which is unsuitable for observing cellular morphology (reviewed by Saura, 2007). Morphology and F-actin localization of fibroblast-like astrocytes are almost the same as that of fibroblasts, which have polygonal shape and thick stress fibers containing F-actin (Güler et al., 2021). Indeed, transcriptome analysis indicated that fibroblast-like astrocytes highly express hundreds of genes that are not normally expressed in astrocytes in vivo (Foo et al., 2011). Some studies indicated that PAPs of fibroblast-like astrocytes contained actin-binding proteins such as ezrin and profilin (details are described in discussion) (Lavialle et al., 2011; Molotkov et al., 2013). However, this was in reference to the thin membrane protrusions directly extending from the cell body as PAPs.

Thus, the factors affecting astrocyte morphology have been actively analyzed in tissues and animal models. Knockout or transgenic mice, in which astrocytes change their morphology, also provide information about specific genes. In living animals and tissues, adeno-associated viruses (AAVs) with astrocyte-specific promoters have been used as a powerful tool for transfecting genes into astrocytes (reviewed by Shigetomi et al., 2016). For example, AAVs utilizing shRNA of TrkB.T1, a brain-derived neurotrophic factor (BDNF), reduced the volume and complexity of astrocytes in vivo (Holt et al., 2019). Recent RNA-seq findings have also provided extensive information. For example, putative actin-interacting proteins FERMT2 (FERM domain containing kindlin) and ezrin have been reported as astrocyte territory size-related modules based on analysis of RNA-seq and astrocyte volume in brain tissues (Endo et al., 2022). For uncovering the molecular basis of nanostructures of astrocytes in tissues and animals, electron microscopy provides high-resolution protein localization; however, the method requires the fixation of tissues. Two-photon microscopy or super-resolution microscopy (reviewed by Arizono et al., 2023) allows the monitoring of protein dynamics in living tissues. However, the methods are costly and furthermore, currently available imaging methods in living tissues are difficult for demonstrating protein dynamics.

Thus, the development of a cell culture system enabling high-resolution 2D observation is anticipated to facilitate knowledge sharing among researchers. Previously, we reported that a culture method of chicken astrocytes in Neurobasal medium containing b-FGF and B27 formed stem processes at low cell density. Lamellipodia and filopodia containing F-actin were observed in the cell periphery and specifically concentrated in stem process tips (Tsukuda et al., 2019). However, localization of cytoskeletal proteins other than actin and GFAP has not been analyzed using immunostaining. Additionally, we have not observed behaviors of proteins because transfection methods that maintain these structures in cultured astrocytes have not been previously developed.

The current study has two main aims. First, we aimed to observe localization of cytoskeletal proteins in detail during the development of our cultured astrocytes. Additionally, we analyzed morphological heterogeneity regarding the number and length of branches, frequency of branching, and morphology of process tips. The second aim was to analyze the behaviors of actin-binding proteins in living astrocytes, whereby we examined various transfection methods (lipofection, electroporation, and lentivirus infection) that maintain stem processes and PAPs at low cell density that do not overlap with other cells. For preparing large quantities of homogeneous cells to examine various transfection conditions, we attempted to culture astrocytes that were directly cryopreserved after dissociation from the forebrain (directly cryopreserved astrocytes). In this study, to evaluate our culture method, we focused on the localization and behavior of two actin-binding proteins, ezrin and lasp-2. Ezrin is an anchor between actin and membrane, and is a well-studied standard PAP marker for analyzing astrocyte morphology (reviewed by Derouiche & Geiger, 2019). The other protein, lasp-2, first identified in our group, has been reported to be localized in filopodia, lamellipodia, and focal adhesions in various types of cultured cells (reviewed by Butt & Raman, 2018). mRNA of lasp-2 and another actin-binding protein, nebulette, are generated from a single gene (LASP2/NEBL) as splice isoforms in vertebrates (Fujita et al., 2022); lasp-2 is highly expressed in the brain and nebulette is specifically expressed in cardiac muscles (Terasaki et al., 2004; Zieseniss et al., 2008). Localization of lasp-2 has not been studied in astrocytes, but high expression in cultured astrocytes was confirmed in our previous study (Tsukuda et al., 2019). Thus, both proteins are used to examine whether localization varies across different actin-containing structures visualized by phalloidin-staining or fluorescent-tagged actin in cultured astrocytes.

The present study basically follows terms defined by Baldwin et al., (2024) (Sup. Fig. 1). Based on recent morphological studies, they classified astrocyte processes in tissues into three categories that convey meaningful information. They suggested ‘branches’ are the major stem processes emanating from the astrocyte soma, ‘branchlets’ are the finer secondary structures emanating from branches, and ‘leaflets’ are the thinner regions of astrocyte branches and branchlets that contact synapses. In the current study, we also use “branches” and “branchletes” to refer to processes more than 10 μm in length observed in phase-contrast images and used “stem process” for branches and branchlets. Additionally, we defined “somal protrusions” and “collateral protrusions” as 5-10 μm protrusions, which are expected to grow into new stem processes. We also newly named “collateral veils” as thin membrane ruffles that are difficult to identify in phase-contrast images but contain microvilli clearly stained with anti-ezrin antibody. Baldwin et al., (2024) noted that truly “peripheral” astrocyte processes do not exist, because fine processes exist throughout astrocyte territories and “perisynaptic” astrocyte processes should be identified in processes specifically enwrapping synapses. However, the term “PAP” is not suitable for our astrocyte culture, since there are few neurons. Instead, we use the term “PAP-like structures” to describe actin-rich structures observed in the peripheral region of astrocytes. Observation of the PAP-like structures in high resolution suggested that they consist of various actin-rich substructures of lamellate processes and filamentous processes, which are actively moving. Moreover, in the inner cell area, stress fiber-like structures and elliptical structures which are considered to cell-substrate adhesions were also observed.

We successfully observed two actin-binding proteins (ezrin and lasp-2) in cultured astrocytes retaining stem processes and PAP-like structures and revealed that they localized in different actin-rich substructures. Findings in the current study indicated that our cultured astrocytes mimic the development of astrocytes, which extend their cell territories to reach target cells, and that the improved protocols offer significant advantages in future astrocyte research to investigate the roles of actin cytoskeleton.

## RESULTS

### Chicken astrocytes offer a suitable cell model that mimics the development and morphology of astrocytes in vivo

We compared the development of direct cryopreserved (DC) astrocytes with freshly harvested (FH) and pre-cultured cryopreserved (PC) astrocytes prepared using our previous methods (Tsukuda et al., 2019). Around 5-10 × 10^5^ cells per 15-day chick embryo forebrain were collected, and DC astrocytes after thawing had a ∼70% recovery rate of ∼70% and ∼70% cell viability; thus, approximately 40–60% of the cryopreserved cells were culturable. As we described previously (Tsukuda et al., 2019), most contaminants, including erythrocytes, cell debris, and extracellular matrix, were removed after the first medium change on the next day. Storage of astrocytes in a deep freezer (-80C) reduced both recovery rate and viability, with lower number of culturable cells (25-40% after 6 months). Liquid nitrogen retained cell viability for at least two years, similar to cells just after cryopreservation.

The increase in coverage of DC astrocytes and PC astrocytes was slower than for FH astrocytes at all cell densities (Fig. S2A). At day 3 of culture, DC astrocytes and FH astrocytes showed similar rates of expression of GFAP as an index of differentiation, and PC astrocytes tended to show rather higher expression (Fig. S2B), which is similar to our previous study. At day 8 of culture, the GFAP-positive rate varied with density and culture method, but all remained above 70% in FH and PC astrocytes and above 60% in DC astrocytes (Fig. S2B). There was no significant difference in morphology of DC astrocytes in phase-contrast images (Fig. S2C), and all DC astrocytes displaying more than 4 branches were GFAP-positive, as reported in FH astrocytes (Tsukuda et al., 2019).

Using staining with anti-GFAP antibody and phalloidin, we confirmed that the morphological differentiation of DC astrocytes (Fig. 1; see also Fig. S1 for terminology) was similar to that of FH astrocytes. When DC cells were cultured at an initial density of 0.1 × 10^5^ cells/cm^2^ (0.1id) for 3 days, short rod-shaped or Y-shaped cells under 100 μm in length were observed (Fig. 1, upper left panel). Some cells had branches with stick-like process tips (>10 µm, arrowhead) and short protrusions directly extended from soma (5-10 µm, square bracket), which we defined as “somal protrusions” and are expected to grow into new branches. About 45% of the cells expressed GFAP (Fig. S2B), but the staining was not filamentous but dotty. On day 5, 0.1id astrocytes generated multiple branches with GFAP staining stronger than on day 3 (Fig.1, upper middle). Some stem processes extended over 50 μm with short protrusions emerging from the sides of stem processes (5-10 µm, square bracket), which were defined as “collateral protrusions” and are expected to grow into new branchlets. Accumulation of actin meshwork was observed in both somal and collateral protrusions, and all protrusions under 20 µm were stained with phalloidin but not anti-GFAP. The length of the cell area (covered with short branches and branchlets) of some astrocytes extended over 100 µm. The width of process tips identified in phase-contrast images varied, and both stick-like process tips (arrowheads, with process width < 2 times the branch/branchlet width) or larger palm-like process tips (arc, with process width > 2 times the branch/branchlet width) were stained with phalloidin.

**Fig. 1.**
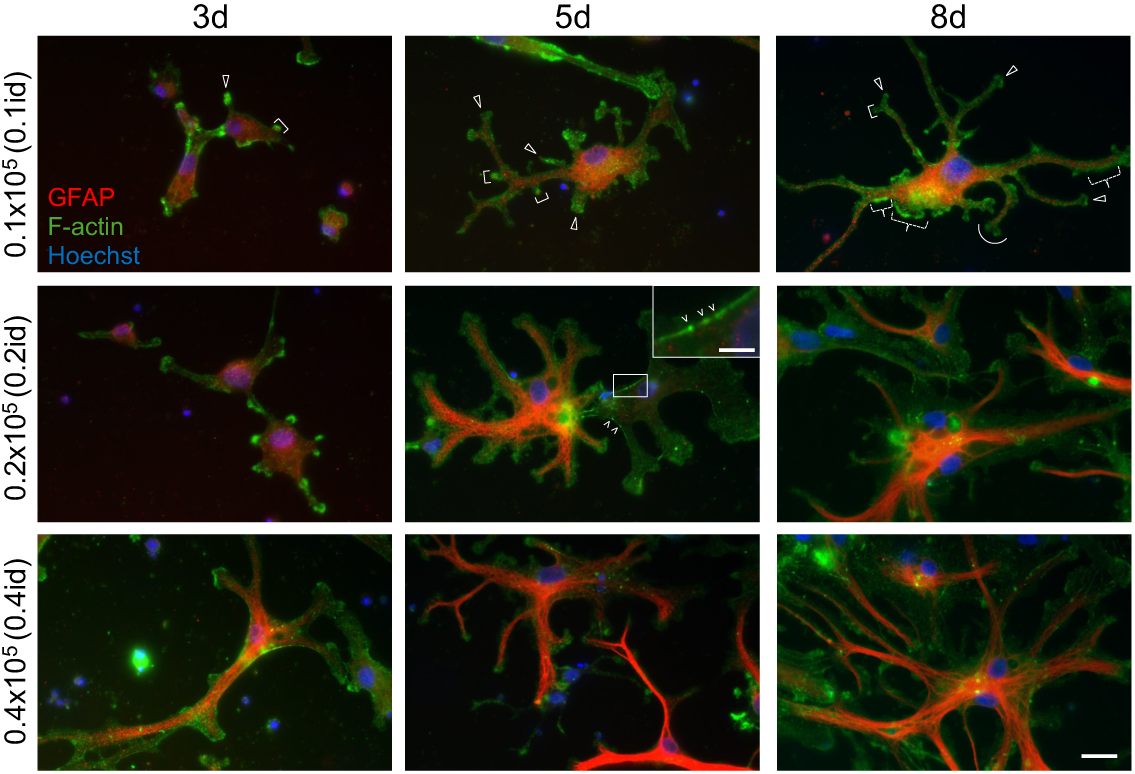
Representative images of astrocytes at different initial cell densities. Representative images of direct cryopreserved astrocytes stained with anti-GFAP antibody (red), phalloidin (green), and Hoechst 33342 (blue) at different initial cell densities after 3, 5, and 8 days. Examples of “protrusions directly emerged from soma”/“collateral protrusions” (square brackets), “collateral veil” (dashed curly brackets), “stick-like process tips” (arrowheads), and “palm-like process tips” (arcs), as described in Fig. S1, are indicated only in the top panels. Wedges indicate irregular and stronger staining of F-actin in stress-fiber-like structures. All figure annotations apply to subsequent figures. Scale bar = 20 μm; areas highlighted by the squares are magnified, with scale bar = 5 μm (same scales apply to fluorescent images in subsequent figures).

On day 8, the coverage area of 0.1id astrocytes reached about 20% (Fig. S2A); most of the astrocytes were observed as isolated cells, and palm-like process tips (arcs) were observed more frequently (Fig. 1, upper right). At this later stage, F-actin staining increased in the peripheral region of astrocytes and we noted these structures as “PAP-like structures” in the present study. The PAP-like structures contained both lamellate processes and filamentous processes intensely stained with phalloidin and, following observations in living astrocytes, confirmed their active movement (see Fig. 4 and 5). There are at least two types of lamellate processes; one type is “collateral veils”, which we named in the current study as thin membrane ruffles difficult to identify in phase-contrast images (see Fig. 2) but contain microvillus protrusions clearly stained with phalloidin (dashed curly brackets). Collateral veils besides stem processes were often observed in cells cultured for over a week (see Fig. 2) and also observed in junctions between stem processes (See Fig. 3A). Another type is lamellipodia mainly observed at the palm-like process tips in phase-contrast images (see Fig. 2). Filamentous processes were difficult to observe in phase-contrast images regardless of their length (described later in Fig. 2D). Long filamentous processes were observed both in stick-like and palm-like process tips, with a higher frequency of the latter. Short filamentous processes directly emerging from stem processes were also observed. Most stem processes over 10 µm were stained with the GFAP antibody and some astrocytes exhibited filamentous GFAP. The length and number of branches and branchlets radially increased, expanding their territories over 200 µm diameter until 7-10 days after plating (see Fig. 2). Their territories seemed to remain stable for at least 4 weeks, although new branching continued slowly.

**Fig. 2.**
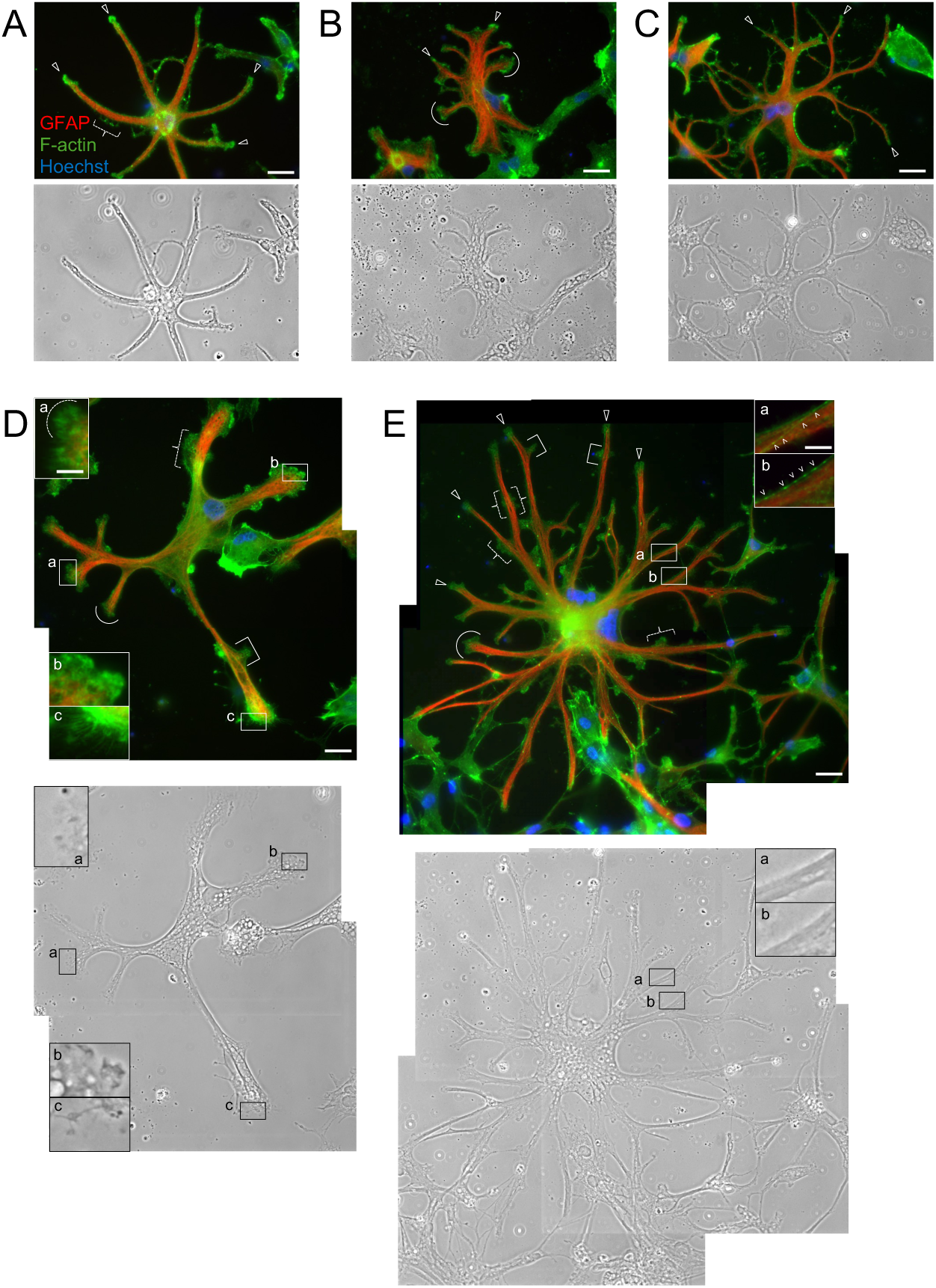
Heterogeneity of cultured astrocytes. Cultured astrocytes with various morphologies. Cells were cultured for 8 days and stained with the same antibodies and reagents as in Fig. 1. (A) Astrocyte displaying thin branches with stick-like process tips (arrowheads) and collateral veils (dashed curly bracket); (B) astrocyte showing thick branches with palm-like process tips (arcs); (C) astrocyte showing multiple branches of various widths with many secondary/tertiary branchlets; (D) astrocyte with variable morphology of process tips; (E) astrocyte with territory of diameter over 100 µm.

**Fig. 3.**
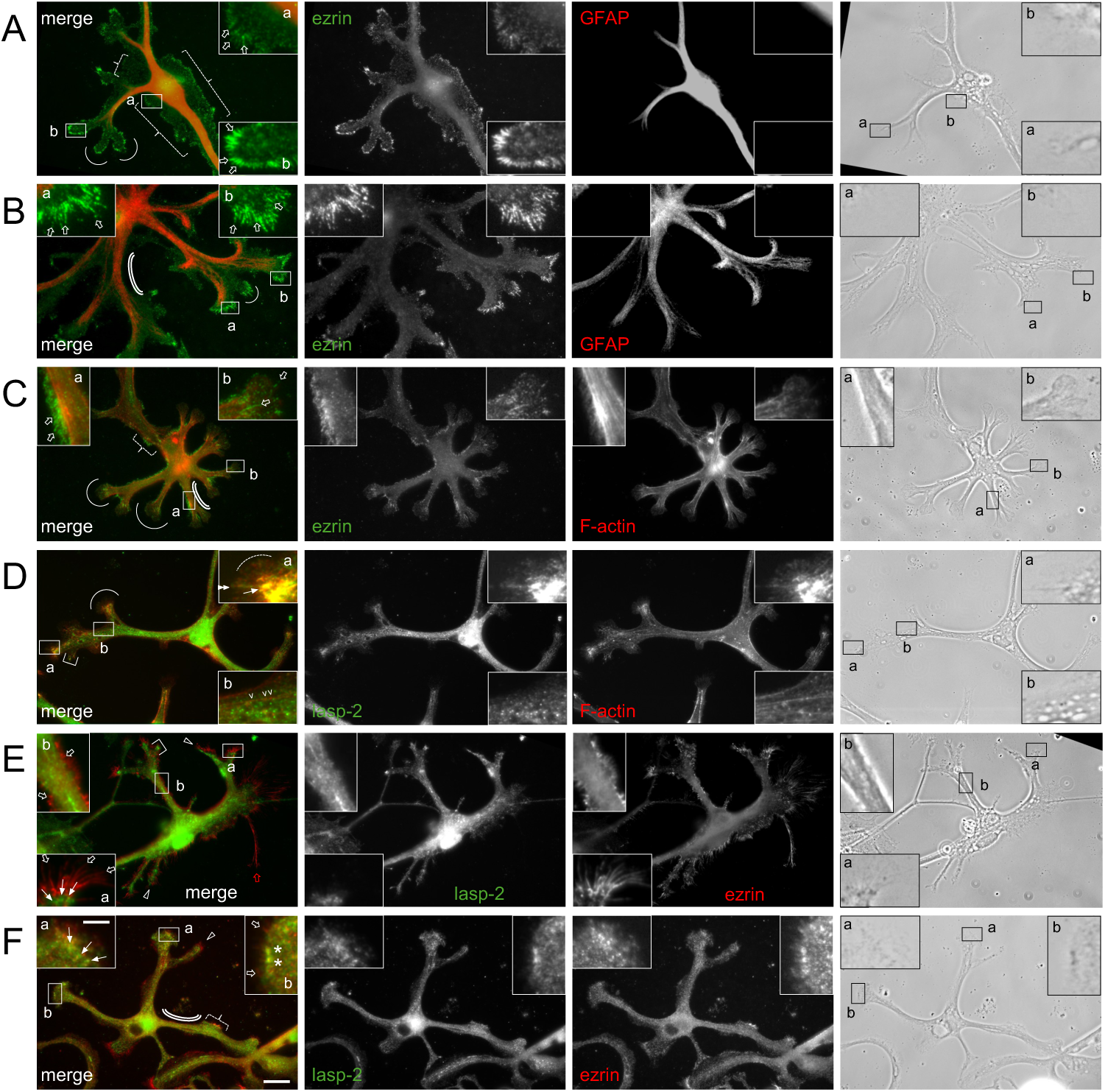
Localization of ezrin and lasp-2 in cultured astrocytes. Representative immunostaining images of (A, B) ezrin (green) and GFAP (red) after 11d; (C) ezrin (green) and F-actin (red) after 10d; (D) lasp-2 (green) and F-actin (red) after 7d; (E, F) lasp-2 (green) and ezrin (red) after 7d. Double arrowheads and dashed arcs indicate filopodia and lamellipodia, respectively. Open arrows indicate ezrin-positive microvilli (this also applies to all subsequent figures). A red open arrow indicates long string-like structures stained with anti-ezrin. Double lines in (B), (C), and (F) indicate examples of areas where anti-ezrin is hardly stained. Arrows in (D), (E), and (F) indicate elliptical structures stained with anti-lasp-2. Asterisks in (F) indicate the inner edge of an example area containing structures where lasp-2 and ezrin colocalized (same applies for Fig 4D’ and 5F’). Brightness of some enlarged images was enhanced to visualize fine structures (same adjustments apply to fluorescent images in subsequent figures).

**Fig. 4.**
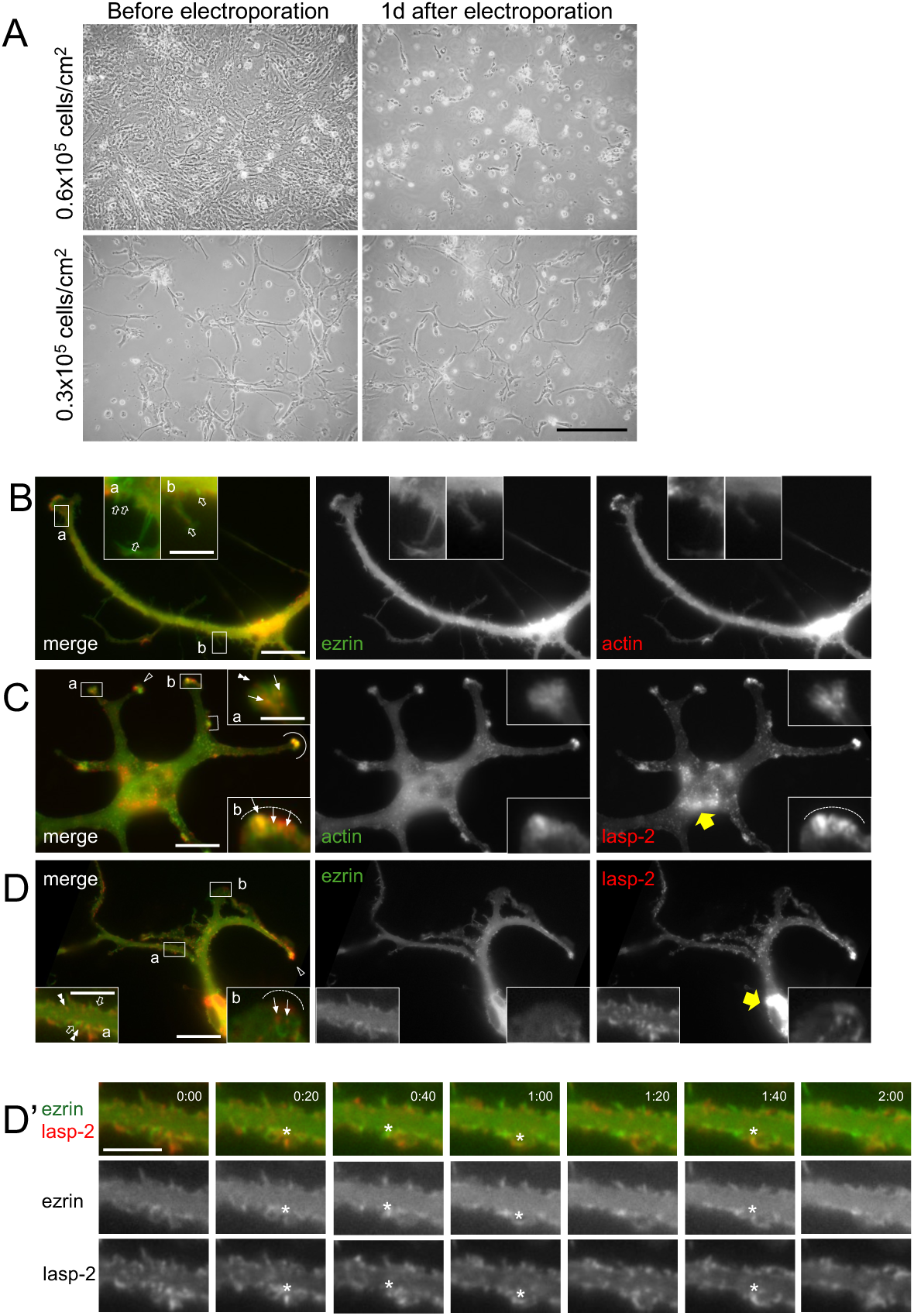
Morphology of astrocytes transfected by electroporation and localization of fluorescent-tagged proteins. (A) Lower magnification view of astrocytes cultured at high density (initial cell density of 0.6 × 10^5^ cells/cm^2^; cultured for 7 days) and low density (initial cell density of 0.3 × 10^5^ cells/cm^2^; cultured for 6 days). Phase contrast images show cells just before trypsin treatment (pre-transfection) and cultured for one day after electroporation. Scale bar = 200 μm. (B-D) Representative images of (B) EGFP-ezrin and mCherry-actin, (C) EGFP-actin and mCherry-lasp-2, and (D) EGFP-ezrin and mCherry-lasp-2 at 2, 3, and 2 days post-transfection, respectively. Yellow arrows in (C) and (D) indicate aggregates of mCherry-tagged protein sometimes observed around the nucleus. See supplementary movie 1 for (D); (D’) kymographs of magnified segment of stem process surface from supplemental movie 1 with a sequence interval of 20 sec. Arrows in (C) and (D) indicate examples of elliptical structures of fluorescent-tagged lasp-2. Asterisks in (D’) indicate examples of structures where lasp-2 and ezrin colocalized.

**Fig. 5.**
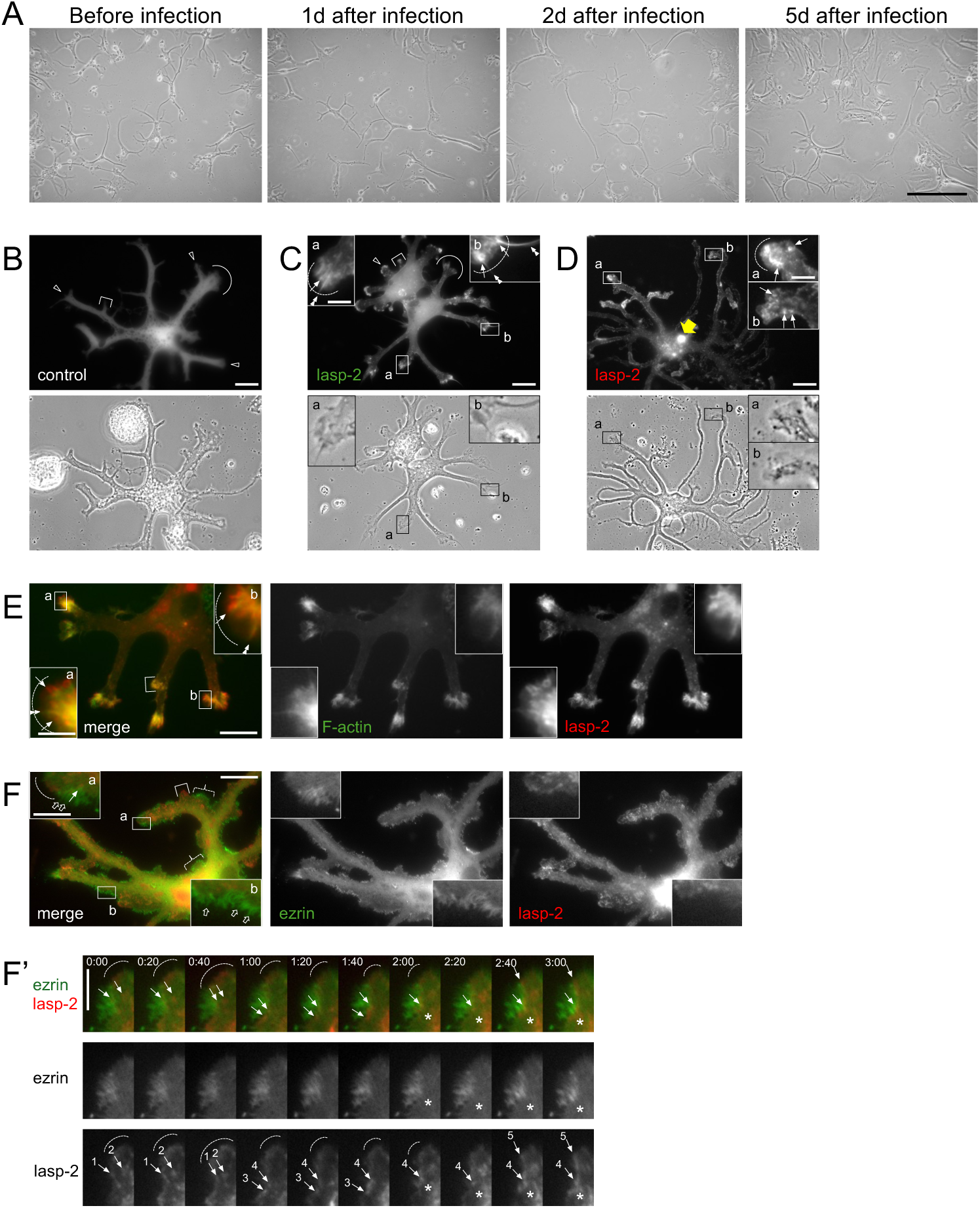
Morphology of astrocytes infected with lentivirus and localization of fluorescent-tagged proteins. (A) Lower magnification view of astrocytes before and after (1d, 2d and 5d) infection. Scale bar = 200 μm. (B-E) Representative images of (B) EGFP, (C) EGFP-lasp-2, (D) mCherry-lasp-2, and (E) mCherry-lasp-2 and added Sara Fluor 497 actin probe in astrocytes at 7, 4, 6, and 5 days post-infection with lentivirus vectors, respectively. Open arrows in (D) indicate large aggregates of mCherry-tagged protein sometimes observed around the nucleus. (F) EGFP-ezrin and mCherry-lasp-2 at 2 days post-infection. See supplementary movie 2. (F’) Kymographs of magnified segments of process tips from the movie with 20 sec sequence intervals. Arrows indicate examples of elliptical structures of fluorescent-tagged lasp-2 and numbered arrows in (F’) indicate individual elliptical structures in image sequences. Asterisks indicate examples of structures where lasp-2 and ezrin colocalized.

Astrocytes cultured at initial densities of 0.2 × 10^5^ cells/cm^2^ (0.2id) after 3 days (Fig. 1, middle left) showed morphology similar to 0.1id cells after 5 days. On day 5, 0.2id astrocytes differentiated in a similar manner to 0.1id astrocytes after 8 days, and 0.2id astrocytes after 8 days exhibited filamentous GFAP more frequently than 0.1id astrocytes. Most of the 0.2id astrocytes formed exposed stem processes and some astrocytes were isolated, which is suitable for observing stem processes. Some astrocytes after 5 days had stress fiber-like structures along stem processes, and irregular and stronger staining of F-actin along these stress fiber-like structures was observed (Fig. 1, wedges in middle center, also see Fig. 2E and 3D). Astrocytes cultured at 0.4 × 10^5^ cells/cm^2^ initial densities (0.4id) differentiated faster than 0.2id astrocytes, but coverage area reached 70% on day 8 (Fig. S2A; Fig.1, lower right). As described previously (Tsukuda et al., 2019), contaminated epithelial-like cells, fibroblast-like cells, and neurons were easily identified by their lack of GFAP staining and their strong polygonal phalloidin staining or long axons.

The morphology of GFAP-positive 0.1id and 0.2id astrocytes cultured for 8 days, which were considered to have finished territory expansion, were observed in detail to analyze the numbers and lengths of branches and branchlets, as well as the width of branches and morphology of process tips (Fig. 2). Some astrocytes had thin and long branches over 50 µm without branchlets and all process tips were stick-like (arrowheads) (Fig. 2A), whereas some other astrocytes had thick branches less than 30 µm without branchlets and almost all process tips were palm-like (Fig. 2B). Most astrocytes had multiple branches with various widths and formed many secondary/tertiary branchlets (Fig. 2C). Palm-like structures formed in the process tips also varied in shape. Most tips had wavy peripherals accompanied with lamellipodia and filamentous processes (Fig. 2D, enlarged image a), although some tips were bumpy without filamentous processes (enlarged image b), and some tips had large bulges with long filamentous processes (enlarged image c). Process tip widths varied from 5 µm to 30 µm. Collateral veils were observed in various types of astrocytes (Fig. 2A, 2D, and 2E). In some astrocytes, the territories of the cells, including stem processes and cell bodies expanded around 300 µm in diameter, and branch numbers ranged over 10 (Fig. 2E).

### Immunostaining of ezrin and lasp-2 showed different patterns in astrocytes

To visualize the localization of actin-binding proteins in actin-rich structures of astrocytes, we compared the localization of two actin-binding proteins, ezrin and lasp-2 using specific antibodies. Microvillus protrusions at the edges of collateral veils were always stained with anti-ezrin (Fig. 3A, 3C, and 3F, dashed curly bracket) and enlarged image confirmed ezrin localization in microvilli (Fig. 3A, open arrows in enlarged image a), but the frequency of ezrin-positive protrusions in other structures varied among cells. In some cells, ezrin-positive microvilli (indicated with open arrows) were observed as dense brush-like structures radially emerging from the process tips (Fig. 3A, enlarged image b; Fig. 3B, enlarged image b; Fig. 3E, enlarged image a) but in some other cells, microvilli were sparsely distributed around the process tips (open arrowheads in Fig. 3C, in enlarged image b; 3F, enlarged image b). Microvilli that directly emerged from stem processes were dense in specific areas (Fig. 3C, enlarged image a; 3E, enlarged image b) but sparse in other areas of the stem processes (Fig. 3B, 3C, and 3F, double lines). The diameter of microvilli seemed to be uniform in all areas but the length varied up to 50 µm (e.g. Fig. 3E, long string-like structures indicated by a red open arrow). Phalloidin staining to visualize actin filaments moderately overlapped with ezrin staining; this may be due to fewer filaments in microvilli than filopodia (Fig. 3C).

Lasp-2 was concentrated with F-actin in lamellipodia and filopodia at most process tips (Fig. 3D, dashed arcs and double arrowheads, respectively, in enlarged image a) and their colocalization was prominent in live cell imaging (described later in Fig. 4C and 5E). Lasp-2 was also observed in collateral protrusions with F-actin, which are expected to be new branches or branchlets (Fig. 3D, square bracket). In magnified images of the process tips, stronger lasp-2 and F-actin colocalization at the base (inner part of tips) of lamellipodia and filopodia was observed as elliptical structures of 1-2 µm along the direction of branch extensions (Fig. 3D, arrow in enlarged image a). Colocalization patterns of lasp-2 and F-actin in these elliptical structures were similar among astrocytes regardless of the length and width of their stem processes. In contrast, lasp-2 did not colocalize with F-actin in all stress fiber-like structures (Fig. 3D, wedges in enlarged image b). There was no lasp-2 localization in ezrin-positive microvilli of process tips, stem processes, and collateral veils, and vice versa: ezrin was not localized in collateral protrusions, lamellipodia, filopodia, and elliptical structures where lasp-2 was concentrated (Fig. 3E and 3F). In a small part of the process tips, ezrin and lasp-2 showed colocalization as small dotty structures in the peripheral region of the tips (Fig. 3F, asterisks in enlarged image b). Some process tips had neither microvilli containing ezrin nor filopodia containing lasp-2 (Fig. 3F, enlarged image a).

### Improved method of electroporation can visualize behaviors of actin-binding proteins in living astrocytes

Using DC astrocytes, we examined transfection methods that retain astrocyte morphology at low cell density. Astrocytes cultured in 60-mm dishes at an initial density of 0.2-0.8 × 10^5 cells/cm² for 6-8 days reached 40-80% coverage, yielding 5-10 × 10^5 dissociated cells. Within 2 days of culture after electroporation, most cells had regenerated stem processes and some of the isolated cells were suitable for observation, although some cells tended to form colonies (Fig. 4A). Appropriate number of cells for transfection was about 0.4-0.8 × 10^6^ and in cases where cell number was increased to the extreme (e.g. 2.0 × 10^6^), transfection efficiency remained the same, although cells tended to form more colonies. Post-electroporation seeding cell density of ≤ 0.2 × 10^5^ cells/cm^2^ suppressed recovery of cell morphology and that of ≥ 0.8 × 10^5^ cells/cm^2^ resulted in fewer isolated astrocytes; about 20% of cells seemed to survive. The addition of a conditioned medium, which is thought to contain growth factors secreted by astrocytes, significantly accelerated morphological recovery. Moreover, cell culture area also affected the recovery. We initially attempted to culture electroporated astrocytes only in the small central glass portion of a glass-based dish (35-mm plastic dish with a centered 12 mm cover glass bottom), but found that astrocytes cultured in the entire dish base (including the plastic area) recovered faster. The 40-80% cell coverage before electroporation was adequate for both transfection efficiency and cell morphology, and cell morphology after transfection tended to deteriorate when the coverage was over 80% (Fig. 4A, upper panels).

About 5-10% of the electroporated cells expressed fluorescent-tagged proteins and multiple plasmid vectors seemed to be electroporated simultaneously into the same cells; thus, transfection efficiencies of single plasmid and double plasmids with EGFP and mCherry tags were almost the same. Fluorescence was easily observed 1-4 days after electroporation and began to disappear 7 days after electroporation. We found that shorter culture periods of less than 5 days required more days for formation of the stem process and PAP-like structures, and culture periods of over 9 days suppressed morphology recovery. We confirmed that electroporated cells with ≥ 4 branches were also GFAP-positive regardless of stem process length and process tip width (data not shown). After 4 days of transfection, in cells expressing mCherry-tagged proteins, aggregates of fluorescent proteins were sometimes observed around the nucleus and they became more pronounced with culture length (Fig. 4C and 4D, yellow arrows), although localization of mCherry-tagged proteins in other areas was similar to that of EGFP-tagged proteins. Fluorescent-tagged proteins showed almost similar localization patterns to immunostaining. In some cells transfected with pEGFP-Lifeact, a short peptide that binds to F-actin, relatively longer multiple filopodia and thick lamellipodia were observed in process tips more frequently both by electroporation and lipofection (Fig. S3B, double arrowheads and dashed arcs); thus we use constructs coding actin to compare the localization of ezrin and lasp-2.

Accumulation of EGFP-ezrin in microvilli on the cell surface varied among cells and areas and length of ezrin-containing microvilli at process tips and stem processes also varied in a similar manner to immunostaining results (Fig. 4B, open arrows in enlarged image a and b). Similar to the phalloidin staining, co-transfected mCherry-actin was weakly observed in the microvilli. In contrast, transfected lasp-2 prominently colocalized with actin and accumulated in all process tips as well as in collateral protrusions of various widths (Fig. 4C, square bracket). Magnified images of palm-like process tips indicated that lasp-2 was colocalized with actin in lamellipodia, filopodia, and some elliptical structures in a similar manner to immunostaining (Fig. 4C, double arrowheads, dashed arcs, and arrows in enlarged images a and b). However, accumulation of lasp-2 in filopodia and lamellipodia varied among process tips, similar to immunostaining results, although most process tips had elliptical structures containing lasp-2 (Fig. 4D, double arrowheads and arrows in enlarged images b). Cotransfection of EGFP-ezrin and mCherry-lasp-2 indicated that they were localized in different protrusions of stem processes; EGFP-ezrin was observed in microvilli and mCherry-lasp-2 was observed in different filamentous structures, which thought to be filopodia (Fig. 4D, open arrowheads and double arrows in enlarged images a). Kymographs of stem processes also showed that substructures containing ezrin or lasp-2 actively moved, although colocalization was observed only in limited areas (asterisks in Fig. 4D’).

We also attempted lipofection of astrocytes using fluorescent-tagged plasmids, but most astrocytes dissociated from the culture dishes after following the manufacturer’s protocol in both one-step and two-step transfection.. Reducing the reagents to one-fifth of the protocol (for example, 0.3 μL of transfection reagent for 1.0 × 10^5^ cells) and addition of conditioned medium resulted in generation of stem processes, but most cells were rod-shaped or spherical after one-step transfection (Fig. S3A, upper panel). Two-step transfection generated astrocytes with multiple stem processes more frequently, but it was difficult to observe the localization of fluorescent proteins because over 50% confluency was required for recovery of the stem processes (Fig. S3A, lower panel). The number and length of the stem processes in fluorescent-positive astrocytes were lower than those following electroporation (S3B, left). The transfection efficiency of lipofection was about 5-10%. Seven days of culture post-thawing is appropriate for transfection, and 3 days post-transfection is appropriate for observing fluorescent proteins, with fluorescence beginning to fade after around 1 week post-lipofection.

### Lentivirus vectors allow for long-term observation of actin-binding proteins in astrocytes

In contrast to electroporated astrocytes that were dissociated from dishes, lentivirus infection hardly affected the development of astrocytes cultured on dishes (Fig. 5A). Cell morphology was slightly disrupted up to 2 days post-lentiviral infection and some cell debris was observed post-infection (Fig. 5B, phase contrast image), but gradually recovered after day 3. Infection efficiency was around 10-50%, and this efficiency was similar at cell densities up to 80% and between 3 to 7 days of culturing. Similarly, fluorescence from lentivirus vectors was weak up to day 2, but could be observed from day 3 and even after 28 days of culturing (data not shown). Astrocytes infected with lentiviral vectors expressing EGFP, EGFP-lasp-2, and mCherry-lasp-2 had well-maintained morphology of stem processes and process tips compared with lipofection and electroporation (Fig. 5B-5D). In contrast to electroporation (which incorporates multiple plasmid types simultaneously), in lentiviral infection, cells were independently infected with two lentivirus vectors containing EGFP and mCherry tags, respectively (5-30%); thus, cells infected with both vectors are 1-20%.

Using lentiviral infection, control EGFP was uniformly distributed in the cytoplasm (Fig. 5B). In contrast, lasp-2 was concentrated in somal protrusions as well as process tips, which is similar to the observations from immunostaining and electroporation (Fig. 5C, square bracket). At the process tips, lasp-2 was also concentrated in lamellipodia, filopodia, and small elliptical structures (1-2 μm) at the base of filopodia and lamellipodia (Fig. 5C and 5D, dashed arcs, double arrowheads, and arrows, respectively, in enlarged images a and b). We confirmed that localization and fluorescence maintenance period of the mCherry-lasp-2 lentivirus were the same as those of the EGFP-tagged lentivirus (Fig. 5D) and some aggregates of mCherry-tagged proteins were sometimes observed similar to electroporated cells (Fig. 5D, yellow arrow). Sara Fluor 497 actin-probe, an F-actin–binding small molecule, strongly stained F-actin in filopodia and lamellipodia at process tips and was colocalized with mCherry-lasp-2 (Fig. 5E, double arrowheads and dashed arcs in enlarged images of a and b). Sara Fluor 497 actin-probe also stained elliptical structures where lasp-2 accumulated (Fig. 5E, arrows in enlarged image a).

Live imaging of mCherry-lasp-2 and EGFP-ezrin showed similar localization patterns to electroporated cells (Fig. 5F). Ezrin was mainly localized in microvilli both in stem processes and process tips (Fig. 5F, open arrows in enlarged images a and b), although lasp-2 localized in lamellipodia and elliptical structures at process tips (Fig. 5F, dashed arcs and arrows, respectively, in enlarged images a). Kymographs of process tips showed that structures containing ezrin or lasp-2 were actively moving (Fig 5F’) in a similar manner to the structures observed in stem processes of electroporated cells (Fig 4D’). Kymographs also indicated that elliptical structures containing lasp-2 appeared, moved, and disappeared (Fig. 5F’, arrows with numbers); the lifetime of elliptical structures was estimated to be 40-180 seconds or longer. In some image sequences, ezrin was found to be concentrated in the outer area adjacent to elliptical structures containing lasp-2 and there was some overlap in the tips of these structures (Fig. 5F’, the region between arrow 4 and asterisk).

To observe the behavior of actin-binding proteins during elongation of stem processes, we observed time-lapse imaging of lasp-2, which was concentrated in process tips in immunostained, electroporated, and infected cells. Astrocytes were infected with a lentiviral vector of control EGFP or EGFP-lasp-2 on day 3 of culture and then observed on day 5, when cells are expected to expand their territories. Fluorescent and phase contrast images were acquired every 2 hours using a glass bottom dish with a grid to detect the positions of the cells (Fig. 6, see analysis method in Fig. S4). Stem processes of astrocytes sometimes extended, retracted, and disappeared frequently. Most process tips of extending stem processes had palm-like structures, and the retracting stem processes had stick-like tips (Fig. 6, control, a and b; lasp-2, a). Sometimes, a tip split into two and then fused back (Fig. 6, lasp-2, b and b’). Moreover, some collateral protrusions developed into new branchlets (Fig. 6, control, b’) and some collateral protrusions disappeared (Fig. 6, lasp-2, c). Phase contrast images indicated extension and branching of stem processes occurred despite remaining attached to other cells (Fig. 6, control, b and b’). Compared with the control EGFP, EGFP-lasp-2 was concentrated in the process tips of extending stem processes and newly developed collateral protrusions (Fig. 6, lasp-2, a at 0, 4, 6, 8 h; b at 2, 4, 8 h). Relative to extending stem processes, fluorescence intensity of EGFP-lasp-2 at process tips decreased in retracting stem processes (Fig. 6, lasp-2, a at 2 h; b at 6 h).

**Fig. 6.**
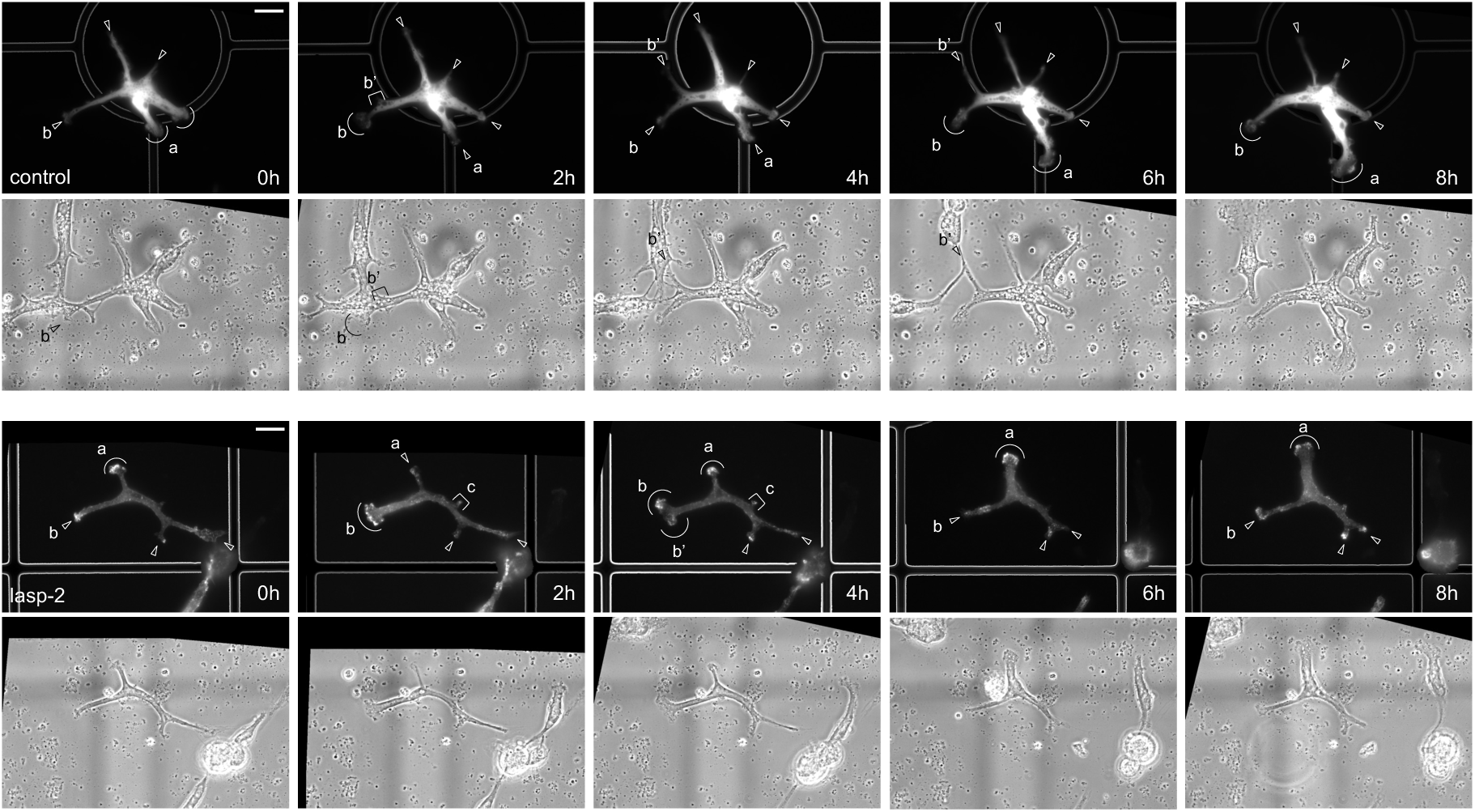
Time-lapse imaging of astrocytes during extension and retraction of stem processes. Time-lapse imaging of astrocytes infected with lentiviral coding with EGFP or EGFP-lasp-2 every 2 hours. Arrowheads, arcs, and square brackets with (a), (b), (b’), and (c) indicate individual process tips in image sequences.

## DISCUSSION

Understanding how neurons and glial cells develop, maintain, and change their elaborate morphology is essential for analyzing neural networks. To achieve this, primary cell culture is a powerful tool, and both a culture method mimicking cell function (including morphology) and a method for transfecting vectors are essential. In primary neurons, established protocols for high-resolution 2D observation of protein localization in neurons revealed the roles of actin-binding proteins in generating growth cones, developing axons, and forming dendrites and spines (reviewed by Lowery & Van Vactor, 2009; Gallo, 2024). Development of the stunning complexity and cell-cell interactions via PAPs of astrocytes also involves extensive cytoskeleton remodeling. However, very little is known about the molecular machinery behind these comprehensive shapeshifts of astrocytes (reviewed by Schiweck et al., 2018). As mentioned in the introduction, a key reason contributing to this limited insight is that the cultured astrocyte models do not accurately recapitulate astrocytes morphogenesis in situ. Recently, culture protocols of rodent astrocytes consisting of several negative and positive immunopanning steps for removing other cell types and collecting astrocytes have been developed, and the purified astrocytes formed stem processes in serum-free medium (Foo, 2013). However, immunopanning requires many kinds of antibodies, of which specificity should be confirmed for the species.

Here, we describe an improved, highly reproducible, and cost-effective protocol for preparation and transfection of chicken astrocytes prepared from embryonic forebrains. From cell coverage area and GFAP staining, we confirmed that directly cryopreserved astrocytes, suitable for accumulating large quantities of homogeneous cells, differentiated similarly to both freshly harvested astrocytes and pre-cultured cryopreserved astrocytes (Fig. 1 and Fig. S1 and S2). Although our culture contained GFAP-negative cells (Fig. S2), we confirmed that cells with 4 or more branches were GFAP-positive astrocytes (Fig. 1 and 2) and used these cells with rich branches for analysis. Directly cryopreserved astrocytes showed heterogeneity in numbers and lengths of branches and branchlets as well as morphology of PAP-like structures (Fig. 2 and 3), suggesting that our cultured astrocytes much more closely resemble in vivo astrocytes. Kálmán (1998) describes the distribution of GFAP-positive astrocytes in chicken brains from hatching to maturity and found that morphology and territories of astrocytes differed among areas and development stages. Although territory size was not explicitly measured, the study’s figures indicate that territories extended up to 400 μm, and our observations that some cultured astrocytes extended their territories to up to 300 µm fit within this range (Fig. 2).

Chickens have been used extensively as research models for developmental studies and as resources for protein purification and primary cultured cells. Low costs, accessibility, ease of manipulation, availability of detailed descriptions of morphogenesis and whole genome sequences (Smith et al., 2023), and recently developed genetic engineering techniques (Panda & McGrew, 2022) have made chickens a robust model for developmental research. Many actin-binding proteins, including lasp-2 analyzed in our study, have been purified from chicken tissues and their amino acid sequences show high similarity to those of mammals. As an example of lasp-2, our previous research indicated that amino acid similarity between chickens and humans is over 90% (Fujita et al., 2022). Moreover, dating as far back as decades earlier, the function of actin-binding proteins have been analyzed in chicken primary fibroblasts, myoblasts, cardiocytes, and sensory neurons, of which morphology and motility were confirmed to mimic cells in vivo (Geiger, 1979; Endo et al., 2007; Zieseniss et al., 2008). Due to the adequate reproduction of in vivo astrocyte development features, such as stem process extension and formation of process tips and collateral veils with microvilli containing ezrin (a well-established PAP marker of astrocytes in tissues) (Fig. 3-6), our cultured chicken astrocytes provide a vital resource for neuroscience research.

Our cultured cells mimicking astrocyte morphology provided high-resolution 2D observation of stem processes and PAP-like structures, albeit with lower stem process complexity. This simpler complexity makes it suitable for analyzing how stem processes extend their territories and how PAPs contribute to stem process elongation and cell-cell interactions. Additionally, measuring cell area and the number and length of stem processes in 2D systems are much easier than in 3D systems. Thus, our system is suitable for analyzing the effects of knockdown or overexpression of proteins, as well as effects of drugs and growth factors in vitro. Moreover, 2D culture systems are applicable for analysis of cell-substrate structures in high resolution using e.g. total internal reflection fluorescence microscopy (TIRFM) and interferometric photoactivated localization microscopy (iPALM) (reviewed by Kanchanawong & Calderwood, 2023). Recently, it has been revealed that the cell-cell adhesion between neurons and astrocytes depends both on direct binding via cell-adhesion molecules (CAMs) and indirect binding via extracellular matrix (reviewed by Saint-Martin & Goda, 2022), and our cultured astrocytes could be applied to co-culture systems to observe such interactions because chicken forebrain neurons are readily prepared from embryos of early development stages (Pettmann et al., 1979).

We also consider that previous studies of actin cytoskeleton utilizing cultured fibroblast-like astrocytes could adapt our newly established protocols. As an example, a study in such cells indicated that formation of PAPs (referred to as thin membrane protrusions directly emerging from the cell body) required ezrin in the membrane/cytoskeleton-bound form (Lavialle et al., 2011). Another study indicated that overexpression of mutant profilin affected the number and length of PAPs, which are also observed as thin membrane protrusions of fibroblast-like astrocytes (Molotkov et al., 2013). Localization of paxillin in focal adhesions of such astrocytes have also been reported (Güler et al., 2023). However, there are no prior reports observing the localization of the above proteins in PAPs of astrocytes in the brain, except ezrin, and it is difficult to directly compare protein localization between fibroblast-like astrocytes (without stem processes) in culture and astrocytes (with multiple stem processes) in the brain. The current study showed highly similar ezrin localization between our cultured cells and astrocytes in the brain as well as morphological similarity. Our protocols can elucidate knowledge gaps of the function of cytoskeletal proteins between cultured cells and tissues.

In the current study using vectors coding actin-binding proteins (ezrin and lasp-2) to observe actin cytoskeleton in living astrocytes, we optimized transfection protocols and successfully improved two of three protocols (electroporation and lentivirus infection) that retained astrocyte morphology at low cell density (Fig. 4 and 5). Electroporation is easy and suitable for high-throughput analysis of localization of multiple proteins identified by large-scale analysis such as RNA-seq and biotin identification (Bio-ID). However, the number of completely isolated cells is lower relative to lentiviral infection, and fluorescence was observed for only one week (Fig. 4). In contrast, lentivirus infection is a more complex approach but the integration of viral genome into the host cell genome allows for long-term gene expression. Indeed, our lentiviral vectors adequately retained cell morphology and fluorescence for at least 4 weeks (Fig. 5); thus, this method will be beneficial for analyzing behaviour of protein during astrocyte development in detail. We observed that Lifeact affected filopodia length and lamellipodia thickness (Fig. S3B), and recent reports have raised concerns about Lifeact-associated artefacts in some cell types (Flores et al., 2019); thus, we used EGFP/mCherry-actin for visualizing the whole actin cytoskeleton (G-actin and F-actin)(Fig. 4B and 4C). Additionally, we confirmed that chemically synthesized F-actin probes easily visualize actin cytoskeleton with actin-binding proteins (Fig. 5E). Ezrin and lasp-2 localization in transfected cells were similar in both live imaging and immunostaining (Fig. 3 and 4); thus, we conclude that our improved transfection protocols are suitable for investigating the roles of actin-binding proteins in astrocytes.

In this study, various actin-containing structures in the cell periphery and inner region of cells were observed in our cultured astrocytes (Schematic illustration shown in Fig. S1). Actin meshwork always accumulated in both somal protrusions and collateral protrusions emerged from branches or branchlets, indicating that the formation of stem processes and branching depends on actin cytoskeleton (Fig. 1). Accumulation of F-actin was observed in process tips, which varied in shape, indicating that elongation of stem processes also depends on actin cytoskeleton (Fig. 2D). Surrounding the cell periphery, six types of “PAP-like structures” (classified in Fig. S1B) were observed as actively moving structures, suggesting that cultured astrocytes have active ruffles similar to those observed in vivo. Thin stress fiber-like structures (indicated with wedges in Fig. 1-3) were markedly different from the thick stress-fiber-like structures commonly observed in fibroblast-like astrocytes that contained a high activation state of myosin-II (John, 2004). However, extending stem processes may depend on the tension of stress fiber-like structures because these structures were observed along stem processes. Some cell types have stress fibers with discontinuous localization of mechanosensing proteins, such as zyxin and nonmuscle myosin, that make up the contractile apparatus (reviewed by Smith et al., 2014). The irregular and stronger staining of F-actin along stress fiber-like structures may be related to the irregular localization of such proteins, and thus, future research is needed. We confirmed localization of ezrin in microvilli in three types of PAP-like structures (summary in Fig. S1): (i) at the edge of collateral veils (one of the types of PAP-like lamellate structures); (ii) directly emerged from stem processes; and (iii) radially emerged from the process tips (Fig. 3A, 3B, and 3E). These structures resemble structures observed in PAPs in tissues, because the diameter of microvilli seemed to be uniform across both tissues and our cultured cells, based on the similar ezrin staining pattern (Lavialle et al., 2012). The microvilli of intestinal brush borders, which have a uniform number of actin filaments of around 30-40 (Ohta et al., 2012), also have a uniform diameter; thus, the actin filament number in astrocyte microvilli may be regulated in the same manner. These protrusions were difficult to identify in phase-contrast images of our cultured astrocytes, but anti-ezrin staining indicated that astrocytes may interact with other cells over a wide area including stem processes and process tips (Fig. 3A). The frequency of ezrin-containing microvilli varied among cells and areas in our cultured astrocytes, suggesting that ezrin locally controls cell-cell interactions in vivo. In contrast, lasp-2 is always localized in the same structures, as we discuss next.

In the present study, we also revealed that our cultured astrocytes formed three additional types of PAP-like structures, (iv) filopodia directly emerged from stem processes, (v) lamellipodia at process tips, and (vi) filopodia at process tips (in Fig. S1), that have various widths and contain lasp-2 but not ezrin. In addition to filopodia and lamellipodia in process tips, the anti-lasp-2 antibody always stained somal protrusions and collateral protrusions expected to become new branches and branchlets, indicating that lasp-2 always binds to F-actin in these structures (Fig. 3D-F). Using Bio-ID AAV vector infected in rat brains to label proteins that localized closely to ezrin, Soto et al. (2023) identified over 100 proteins including the gene product of LASP2/NEBL, which generates lasp-2 and nebulette. Since mRNA of nebulette is mainly expressed in cardiac muscles, the gene product is considered to be lasp-2. They also indicated that staining of brain sections expressing EGFP-ezrin using anti-nebulette C-terminal antibody (with shared sequence with lasp-2) showed colocalization within the astrocyte territory. However, in our study, lasp-2 did not localize in ezrin-positive structures and, vice versa, ezrin hardly localized in lasp-2-positive structures both in immunostaining and live imaging (Fig. 3E, 3F, 4D, and 5F, summarized in Table 1). It is known that conversion of lamellipodia and filopodia occurs frequently (reviewed by Svitkina, 2018). In contrast, microvilli, which have mainly been investigated in stereocilia of hair cells and brush borders of intestinal epithelial cells, have been observed as independent structures, with ezrin localized exclusively in these protrusions (reviewed by Ansel et al., 2024). Thus, it is reasonable to consider that structures containing ezrin or lasp-2 exist independently. Combining the observations in whole cell images of Soto et al. (2023) with our observations under high resolution, it is considered that PAPs in vivo also contain various actin substructures containing either ezrin or lasp-2 surrounding the cell surface (summarized in Fig. S4), and they likely have different roles in regulating actin cytoskeleton.

Accumulation of lasp-2 in the process tips of extending stem processes and newly developed somal protrusions and collateral protrusions in cultured astrocytes suggests that lasp-2 mainly controls the formation and extension of stem processes forming lamellipodia, filopodia, and cell-substrate adhesions (Fig. 3D, 3E, 4C, 5C, 5E, and 6). On the other hand, varied accumulation of ezrin in collateral veils, process tips, and stem processes suggests that ezrin locally controls cell-cell interaction via microvilli (Fig. 2A, 3B, 3E, 3F, 4D, and 5F). This idea is supported by a recent report that astrocytes of nervous system-specific ezrin knockout mice had reduced volume but satellite morphology (Schacke et al., 2022). Observations that astrocyte-specific depletion of ezrin resulted in shorter astrocyte leaflets and reduced astrocyte contact with the synaptic cleft (Badia-Soteras, 2023) also support this idea. Overlapping of ezrin and lasp-2 colocalized in restricted areas of process tips (Fig. 3F and 5F’) may indicate that some ezrin-containing microvilli were extending from lasp-2-containing lamellipodia and they cooperate in extending stem processes. To understand the function of lasp-2 in the elongation and cell-substrate adhesions of stem processes in more detail, it is necessary in future to suppress lasp-2 expression or transfect deletion constructs of lasp-2 (lacking actin-binding regions or adhesion sites to components of focal adhesions), which we developed and investigated previously in cell lines (Nakagawa et al., 2009).

Additionally, the elliptical structures containing lasp-2 and F-actin are observed both in immunostaining (arrows in Fig. 3D) and live imaging (arrows in Fig. 4C and 5E), might be cell-substrate adhesions, because lasp-2 has been reported to be localized in focal adhesions, an integrin-mediated cell-substrate adhesion complex widely distributed in cell lines (Nakagawa et al., 2009; Bliss et al., 2013). The lifetime of some elliptical structures observed in the current study was shorter (Fig. 5F’, within a minute) than that of common focal adhesions (over minutes). Still, several types of cell-substrate adhesions linked to actin cytoskeleton, with various lifetimes, have been reported (reviewed by Kanchanawong & Calderwood, 2023), and whether any components of the adhesions reported in other types of cells show colocalization with lasp-2 in the elliptical structures should be investigated.

Live cell imaging of astrocytes transfected with fluorescent-tagged vectors expressing actin, ezrin, and lasp-2 revealed that filopodia, lamellipodia, and microvilli are actively moving structures (Fig. 4, 5, Supplemental Movie 1 and 2). We found that free stem processes of astrocytes sometimes extended, retracted, and disappeared frequently (Fig. 6). Axonal elongation of cultured neurons commonly continues until growth cones come in contact with other cells that cause repulsion or synapse formation depending on cell-cell interactions, so the movement of stem processes are somewhat different from neurons. Extension of stem processes of astrocytes is also different, whereby extensions that remain attached to other cells are continuous. We observed lasp-2 concentrated in extending process tips and dissociated from retracting tips (Fig. 6). These observations support the idea that lasp-2 controls the stem process extension of astrocytes through filopodia and lamellipodia motility and adhesion to the extracellular matrix.

The morphology of palm-like process tips with highly motile lamellipodia and filopodia, as well as the concentration of actin and actin-binding proteins (ezrin and lasp-2) observed in this study, are similar to those of neuronal growth cones (Fig. 4C, D, and 5B-E) although extending and retracting behaviors are different (Fig. 6). Growth cone-like structures containing actin-binding proteins have been observed in process tips of oligodendrocytes (reviewed by Thomason et al., 2020) and thus, glial cells may extend their processes during development and form complex neural networks. In neurons, several actin-binding proteins were reported to be responsible for neuronal migration disorders (reviewed by Liu, 2011). Recent studies suggested the involvement of astrocytes in actin-based morphology as described in the introduction, as well as of morphologically abnormal astrocytes in brain disease pathogenesis (Endo et al., 2022; Soto 2024). Additionally, the disruption of astrocyte-neuron interactions in synapses and astrocyte-endothelial cell interactions in the blood-brain barrier have been reported in some diseases (reviewed by Blanco-Suárez et al., 2017). Thus, analyzing the functions of numerous actin-binding proteins that play roles in stem process extension, a process including branch formation, cell-substrate adhesion and cell-cell adhesions, in astrocytes may lead to restoring such abnormalities and partly ameliorating the disease. The methods developed in the present study offer significant advantages for filling in the knowledge gaps regarding protein activities in vitro and the roles of their coding genes analyzed in tissues and animal models, thereby contributing to future astrocyte research.

## MATERIALS AND METHODS

### Antibodies

For immunostaining, Cy3-labeled anti-GFAP mouse monoclonal antibody (clone G-A-5, 1:4000, Sigma-Aldrich, St. Louis, MO, USA), anti-lasp-2 specific rabbit polyclonal antibody immunoabsorbed with lasp-1 (1:5; Terasaki et al., 2004), and anti-ezrin mouse monoclonal antibody (clone 3C12, 1:50, Santa Cruz Biotechnology, Dallas, TX, USA) were used for primary antibodies. Alexa Fluor 488-labeled and Alexa Fluor 546-labeled anti-rabbit and anti-mouse IgG antibodies (1:500 or 1:1000, Thermo Fisher, Waltham, MA, USA) were used for secondary antibodies. Alexa Fluor 488-labeled and rhodamine-labeled phalloidin (0.066 μM, Thermo Fisher) were used for F-actin staining. Hoechst 33342 (0.1 μg/mL, Sigma-Aldrich) was used for nuclear staining.

### Fluorescent protein expression vectors

pEGFP vectors coding chicken lasp-2 were prepared as described previously (Terasaki et al., 2004). pEGFP-lifeact was prepared with the insertion of Lifeact oligo DNA, a 17 amino acid peptide of yeast actin binding protein ABP140 (Riedl, 2008). pEGFP-human beta actin was purchased from Clontech (Mountain View, CA, USA). pEGFP vector coding human ezrin were purchased from Addgene (cat. no. 20680, MA, USA). To generate pmCherry-lasp-2, BamHI-EcoRI fragments of pEGFP-lasp-2 were subcloned into the BamHI and EcoRI sites of pmCherry-C1 vector (Clontech). To generate pmCherry-actin, the XhoI-BamHI fragment of pEGFP-actin was subcloned into the XhoI and BamHI site of pmCherry-C1 vector. All plasmids for electroporation and lipofection were prepared with the Nucleobond Xtra Midi EF (Macherey-Nagel, Düren, Germany).

To generate lentivirus vectors expressing EGFP-lasp-2 and mCherry-lasp-2, the region coding fluorescent-tagged proteins were amplified by PCR using PrimeSTAR Max (Takara, Shiga, Japan), following manufacturer’s protocol. The forward primer to amplify each protein was agaagacaccgactctagaggatccCTACCGGTCGCCACCATG (lowercase indicates primer region for subcloning into the BamHI site and uppercase indicates primer region for connecting the region from CMV promoter to the first methionine of EGFP and mCherry). Reverse primers to amplify cDNA of each protein were as follows: EGFP-lasp-2 and mCherry-lasp-2: tgtaatccagaggttgattgtcgacTGGGGAATTCGGACTAGAGAC (lowercase indicates primers for subcloning into the SalI site and uppercase indicates primers for stop codon of lasp-2). The regions coding EGFP of the pLenti CMV EGFP Puro (658-5) vector (Addgene cat. no. 17448) were removed using BamHI and SalI and replaced with PCR products using NEBuilder HiFi DNA Assembly (New England BioLabs, MA, USA). To generate a lentivirus vector expressing EGFP-ezrin, the region coding fluorescent-tagged proteins were amplified by PCR using the forward primer agaagacaccgactctagaggatccAATTCCCGAAAATGCCGAAACC (uppercase indicates primer region for connecting the region from CMV promoter to the first methionine of ezrin). Reverse primers to amplify cDNA of each protein were as follows: tgtaatccagaggttgattgtcgacTCGCGGCCGCTTTACTTG (uppercase indicates primers for stop codon of EGFP).

### Preparation of chick forebrain astrocytes

Astrocytes were prepared from 15-day-old embryonic chick forebrains, following our previous study (Tsukuda et al., 2019). Briefly, cerebral meninges and superficial blood vessels were removed in phosphate-buffered saline (PBS), and cells were dissociated with 0.1% trypsin-EDTA (WAKO, Tokyo, Japan) for 15 min. Dulbecco’s Modified Eagle Medium (DMEM) containing 10% FBS or calf serum (CS) was added to inactivate trypsin, and cells were then filtered through Kimwipes (Kimberly-Clark, Irving, TX, USA) in a Swinnex filter holder (Merck Millipore, Burlington, MA, USA).Dissociated cells were washed three times by DMEM containing 10% FBS or CS. After the final wash, total cell number in the medium was counted and precipitated cells were dissolved in Cell Banker 1 plus (ZENOAQ, Tokyo, Japan) at 3.2 × 10^6^ cells/mL and stored in liquid nitrogen until use.

### Cell culture

FH cells and cells cultured for 2–3 days and then cryopreserved (pre-cultured cryopreserved astrocytes) were cultured as previously described (Tsukuda et al., 2019); see detailed protocol. For the culture of DC astrocytes, cells in cryotubes were thawed using water at 37°C and washed with DMEM twice. 0.2% Trypan Blue Solution (Sigma-Aldrich) was used as a cell stain to assess cell viability. Cells for immunostaining were plated on a 35-mm culture dish with circular coverslips of 15 mm diameter (Matsunami, Tokyo, Japan). Cells for electroporation or one-step lipofection were plated on a 35-mm culture dish. Cells for lentivirus infection or two-step lipofection were plated on a 35-mm glass base dish (D11140H, Matsunami). Cells for time-lapse imaging were plated on a 35-mm glass base dish with grid (grid size 150 µm) (IWAKI, Tokyo, Japan). All dishes were coated with 0.08 mg/mL Poly-L-Lysine (Sigma-Aldrich) and cell densities were varied in experiments (described later). The culture medium was the same as in Tsukuda et al. (2019): Neurobasal medium (Thermo Fisher) supplemented with 2% B-27 supplement (Thermo Fisher), 0.5 mM L-glutamine, 25 μM Glutamax (Thermo Fisher), and 5 ng/mL b-FGF (WAKO). On the next day after plating, the medium was replaced completely by fresh medium following vigorous pipetting. Half of the medium was changed approximately twice a week.

Cells were imaged using inverted microscopy (CK40, Olympus, Tokyo, Japan) equipped with a digital camera (L-835, Hozan, Osaka, Japan) at 20× magnification. The percentage of area covered by cells at various proliferation stages were manually determined following our previous study (Tsukuda et al., 2019).

### Immunocytochemistry

Astrocytes cultured on coverslips were fixed with 4% formaldehyde in PBS for 5 min at 37°C, and permeabilized with 0.1% Triton-X100 and 10% goat serum in PBS, for 30 min at room temperature. Nonspecific background staining was reduced with signal enhancer (Thermo Fisher) or Blocking One Histo (Nacalai, Kyoto, Japan). For GFAP staining, the cells were stained with Cy3-labeled anti-GFAP antibody diluted in Signal Enhancer HIKARI (Nacalai) for 2 h at room temperature. For lasp-2 and ezrin staining, the cells were stained with primary antibodies overnight at 4°C, then rinsed with PBS, and stained with secondary antibodies for 1 h at room temperature. For observation of F-actin and nucleus, cells were stained with fluorescent-labeled phalloidin and Hoechst 33342 for 10 min. Finally, the cells were mounted on slide glasses using SlowFade Gold antifade reagent (Thermo Fisher).

Following our previous study (Tsukuda et al., 2019), GFAP expression rates were determined from the number of cells identified by nuclear staining with Hoechst 33342 as well as the number of GFAP-expressing cells identified by anti-GFAP antibody staining. Three randomly selected images from each experiment were analyzed.

### Fluorescence microscopy

Cells were observed using an Axiovert135 inverted fluorescence microscope (Carl Zeiss, Oberkochen, Germany) equipped with ×40 phase-contrast objective lens or ×63 DIC lens, fluorescence filters (Zeiss 39, 38HE, and 43HE), and a Coolsnap HQ charge-coupled device (CCD) camera (Teledyne Photometrics, Tucson, AZ, USA) operated by the IPLab software (Scanalytics, Billerica, MA, USA), or an inverted fluorescence microscope (Axiovert135) equipped with fluorescence filters (Zeiss 15 and 17) and a Retiga R1 CCD camera (Teledyne Photometrics) operated by the Micro-Manager.

### Electroporation

Astrocytes cultured at densities of 0.2-0.8 × 10^5^ cells/cm^2^ in 60 mm dishes for 6-8 days (50-80% coverage) were transfected with fluorescent-tagged vectors (pEGFP-actin, pmCherry-actin, pEGFP-lifeact, pEGFP-ezrin, and/or pmCherry-lasp-2) by electroporation using NEPA21 (Nepagene, Chiba, Japan). Medium was collected from the cell culture dishes and stored as conditioned medium. Next, the cells were washed with PBS twice and treated with 0.25% trypsin-EDTA (WAKO) for 2 min. DMEM containing 10% FBS was then added to inactivate trypsin and the cells were detached by pipetting. Cells (0.4-0.8 × 10^6^) were transferred to microtubes, supernatants were carefully removed, and then a mixture of two types of plasmids (1.0-2.5 μg each) and Opti-MEM (Thermo Fisher) was added. The cell mixture was transferred to a NEPA 2 mm cuvette and voltage was applied. The parameters were set to the following values: poring pulse (voltage: 275.0 V; pulse length: 0.3 msec; pulse interval: 50.0 msec; number of pulses: 2; decay rate: 10%; polarity: +), transfer pulse (voltage: 20.0 V; pulse length: 50.0 msec; pulse interval: 50.0 msec; number of pulses: 5; decay rate: 40 %; polarity: +/−). After electroporation, the cells were plated on 35-mm glass base dishes (D11140H, Matsunami) coated with 0.08 mg/mL Poly-L-Lysine at cell densities of 0.2-0.6 × 10^5^ cells/cm^2^ and cultured in Neurobasal medium containing half of the conditioned medium. Transfected cells were observed between 2 to 5 days after electroporation, and the medium was not changed during observation.

### Lipofection

Lipofection was performed according to the manufacturer’s method, except the amounts of transfection reagent and plasmids were adjusted. Astrocytes cultured at densities of 0.2-0.9 × 10^5^ cells/cm^2^ for 2-7 days were transfected with fluorescent-tagged vectors (pEGFP-lifeact and pmCherry) by lipofection using ScreenFect A (WAKO) in two different ways.

One-Step Transfection: Cells were washed with PBS twice and treated with 0.25% trypsin-EDTA (WAKO) for 2 min. DMEM containing 10% FBS was then added to inactivate trypsin and the cells were detached by pipetting. Cells (0.3-0.9 × 10^5^) were transferred to microtubes, suspended in antibiotic-free Neurobasal medium, and a mixture of plasmid and ScreenFect TMA Reagent (WAKO) was added. After the one-step lipofection, the cells were plated on 35-mm glass bottom dishes coated with 0.08 mg/mL Poly-L-Lysine and cultured in antibiotic-free Neurobasal medium.

Two-Step Transfection: Medium was collected from the cell culture dishes and stored as conditioned medium in 4C. Next, cells were washed with PBS once and then a mixture of antibiotic-free medium, plasmid, and ScreenFect TMA Reagent was added. The medium was replaced completely by conditioned medium on the next day after two-step lipofection.

### Lentivirus infection

pLenti-EGFP, pLenti-EGFP-lasp-2, and pLenti-mCherry-lasp-2 were transfected into 293FT cells with psPAX2 (Addgene cat. no. 12260) and pCMV-VSV-G (Addgene cat. no. 8454) using PEI MAX (Polysciences, Warrington, PA, USA) at 40 μg/μL, pH 7.1 to produce virus particles. 293FT cells were cultured in DMEM with 10% FBS, 100 units/mL penicillin, 0.1 μg/μL streptomycin, and 200 mM L-glutamine. 293FT cells cultured in a 75 cm^2^ culture flask for 4 days until 50-90% coverage was replaced with antibiotic-free DMEM and then a mixture of Opti-MEM (Thermo Fisher), PEI MAX, and plasmids was added. After the mixture was reacted for 4-12 h, the medium was replaced with DMEM with antibiotics. After 3 days of culturing, medium (virus particle solution) was collected and filtered using a 0.45 µm pore filter unit (Merck Millipore, Darmstadt, Germany) to avoid contamination of 293FT cells. Virus solutions were stored in cryotubes in liquid nitrogen, thawed under running water, and used for subsequent infections.

Astrocytes cultured for 3 to 7 days were transfected with fluorescent-tagged vectors (pLenti-EGFP, pLenti-EGFP-lasp-2, and pLenti-mCherry-lasp-2) by lentivirus infection. From each cultured astrocyte dish containing 2 mL of medium, 1 mL or the entire volume of medium was removed; then, 2 mL of virus solution in DMEM added 2% B27 and 5 ng/mL bFGF was added. The day after infection, the medium was replaced entirely with Neurobasal medium with antibiotics. For live imaging of F-actin of infected astrocytes, SaraFluor 497 actin probe (HMRef)(Goryo Kayaku, Hokkaido, Japan) was added to the medium at final concentration of 100 nM and observed immediately.

### Time-lapse imaging

Astrocytes cultured on a 35-mm glass dish with 150 µm grid (IWAKI) at a density of 0.2 × 10^5^ cells/cm^2^ for 3 days were transfected with fluorescent-tagged vectors (EGFP and EGFP-lasp-2) by lentivirus infection. Time-lapse images were taken 2 to 4 days after infection. Phase-contrast images and fluorescence images of isolated astrocytes were captured every 2 h up to 8 h. The phase-contrast images of the grid in the same area were captured to monitor the location and elongation of astrocytes. Each binarized fluorescence image of astrocytes was overlaid on a grid image for subtraction of cell area and to create images of the grid excluding cell areas. Next, each subtracted grid image was superimposed onto the respective fluorescence astrocyte image to create fluorescence images with grids. Finally, the fluorescence images with grids were aligned in the same position according to the grid in order to monitor the location and elongation of astrocytes (Fig. S4).

### Statistical analyses

All data are expressed as mean ± standard error of mean (SEM) obtained from at least four independent experiments. “N” refers to the number of independent experiments.

## DETAIL PROTOCOLS

### Materials

Preparation of medium

DMEM with FBS was prepared by mixing the following reagents:

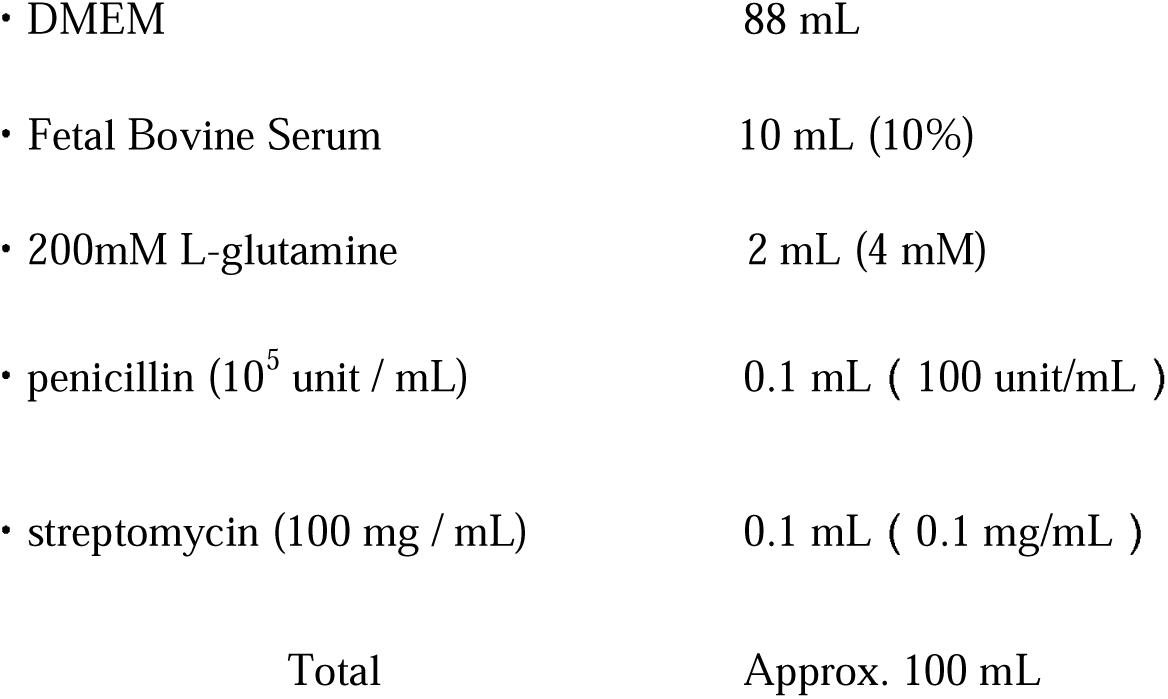

Neurobasal medium with supplements was prepared by mixing the following reagents:

|  |  |
| --- | --- |
| • Neurobasal medium ( Thermo Fisher 21103-049 ) | 49 mL |
| • 200 mM L-glutamine | 0.125 mL ( 0.5 mM ) |
| • 200 mM Glutamax (Thermo Fisher 35050-038) | 6.25 μL ( 25 μM ) |
| • penicillin ( $10^5$ unit/mL ) | 0.05 mL ( 100 unit/mL ) |
| • streptomycin ( 100 mg/mL ) | 0.05 mL ( 0.1 mg/mL ) |
| • B27 supplement ( Thermo Fisher 17504-044 ) | 1 mL ( 2% ) |
| • 10 μg/mL b-FGF (WAKO 064-04541) | 0.025 mL ( 5 ng/mL ) |
| Total | Approx. 50 mL |

B27 supplement and b-FGF should be added just before use.

Preparation of 0.08 mg/mL Poly-L-Lysine (PLL):

- Poly-L-lysine hydrobromide (SIGMA P2636, mol wt 30,000-70,000)

1. Add PBS to the bottle containing PLL to make 0.8 mg/mL solution.
2. Mix well by pipetting and collect it in a 50-mL conical tube.
3. For sterilization, filter the solution into a new tube using a 0.22 µm filter.
4. Aliquot 1 mL portions into 15 mL conical tubes, and store at -20°C.
5. At time of use, prepare a 0.08 mg/mL solution by adding 9 mL of sterile PBS (-) to the aliquot.

□ 0.08 mg/mL PLL can be stored at 4°C for up to 2 weeks.

Preparation of coverslips:

Circular coverslips of 15 mm diameter used in immunostaining were prepared using the following steps.

1. Place 100 pieces of coverslips in a 500-mL glass beaker, add 100 mL of nitric acid, and allow to stand for more than 2 h (stir once every 30 minutes).
2. Remove the nitric acid and rinse the coverslips under running tap water for more than 2 h.
3. Rinse the coverslips with deionized water stirring with tweezers for more than 1 h (change water at least 10 times during the process).
4. Rinse the coverslips 3 times with 50 mL of 100% ethanol.
5. Soak the coverslips overnight in 100% ethanol.
6. Rinse the coverslips once with 50 mL of 100% ethanol.
7. Dry the coverslips in a laminar flow cabinet and store in a case.
8. On the day the cells are cultured, put three coverslips in a 35-mm culture dish and coat with 2 mL of 0.08 mg/mL Poly-L-Lysine for more than 2 h.

Preparation of directly cryopreserved astrocytes:

- Autoclave two Swinnex® filter holders (Merck Millipore, Burlington, MA, USA) inserted with two sheets of Kimwipes (Kimberly-Clark, Irving, TX, USA) and one Swinnex® filter holder with four sheets of Kimwipes.
- Incubate fertilized chicken eggs in the humidified egg incubator at 37.7 °C for 15 days.

1. Set up two 60-mm empty dishes and four 60-mm dishes with PBS.
2. Sterilize eggs (containing chicken embryos) with 70% ethanol. Make a small hole on each of the shells with the back end of tweezers.
3. Gently pick up the chicken embryo with tweezers and transfer to the first empty dish. Repeat for remaining embryos.
4. Decapitate the embryo with the tweezers and transfer only the heads to the second empty dish. Repeat for the remaining embryos.
5. Peel off the hairy skin and remove the skulls to expose the forebrains using the tweezers. Repeat for remaining embryos.
6. Pull out all hemispheres of the forebrains with bent-tip tweezers and transfer them to the first dish with PBS. After gently shaking the hemispheres to remove blood in the first dish with PBS, transfer all hemispheres to the second dish with PBS.
7. In the second dish with PBS, remove the cerebral meninges with blood vessels (thin semi-translucent elastic membranes) using tweezers under a stereomicroscope, and transfer the forebrains to the third dish with PBS.
8. In the third dish containing PBS, tear the hemispheres in half, ensuring that no cerebral meninges remain, and transfer to the fourth dish containing PBS.
9. Transfer all hemispheres to one 50-mL conical tube containing 6 mL of PBS and tear them into 1-2 mm pieces with tweezers.
10. Add 4 mL of 0.25% trypsin-EDTA to make 0.1% trypsin and incubate at 37°C for 15 min. Mix by inversion once every 5 min.
11. Centrifuge the forebrain suspension at 700 rpm for 3 min.
12. Remove the supernatant, add 10 mL of DMEM with FBS, and mix by inversion.
13. Centrifuge the forebrain suspension at 700 rpm for 3 min.
14. Remove the supernatant, add 12 mL of DMEM with FBS, and dissociate cells from forebrain tissues by pipetting 30 times using a 10-mL disposable pipette.
15. Filter the forebrain suspension with a Swinnex® filter holder with two sheets of Kimwipes connected to a 10-mL syringe. Change the holder when the flow decreases (two holders are commonly used to filter suspension from six embryos).
16. Repeat the filter process again but using four sheets of Kimwipes instead.
17. Centrifuge the filtrate at 1200 rpm for 3 min.
18. Remove the supernatant, add 8 mL of DMEM with FBS, and resuspend by pipetting several times.
19. Centrifuge the cell suspension at 1200 rpm for 3 min.
20. Repeat steps 18-19 twice.
21. Remove the supernatant, add 8 mL of Neurobasal medium with supplements, and resuspend by pipetting.
22. Count the cell number with a hemocytometer.
23. Centrifuge the cell suspension at 1000 rpm for 5 min.
24. Remove the supernatant and resuspend the cells in Cell Banker 1 plus at 32 × 10^5^ cells/mL.
25. Dispense the cells in cryotubes at 8, 16, or 32 × 10^5^ cells/tube and store in liquid nitrogen until use.

Thawing and culture of directly cryopreserved astrocytes:

1. Coat the culture dishes or culture dishes with coverslips (for immunostaining) with 2 mL of 0.08 mg/mL Poly-L-Lysine for more than 2 h.
2. Wash the dishes 2 times with PBS. Keep the dish in a 37°C incubator.
3. Defrost cells in cryotubes using a beaker of 37°C water.
4. Transfer thawed cells (in Cell Banker 1 plus) to a 15 mL tube containing DMEM (10 times the volume of Cell Banker 1 plus).
5. Centrifuge the cell suspension at 1000 rpm for 5 min.
6. Remove the supernatant, add 2 mL of DMEM with FBS, and resuspend by pipetting.
7. Centrifuge the cell suspension at 1000 rpm for 5 min.
8. Remove the supernatant, add 1 mL of Neurobasal medium with supplements, and resuspend by pipetting.
9. Count the cell number with a hemocytometer.
10. Adjust the cell suspension to the desired cell density using Neurobasal medium with supplements and place on cell culturing dishes (2 mL for 35-mm dish and 3 mL for 60-mm dish).
11. The next day, to dissociate contaminants (dead cells, erythrocytes, and extracellular matrix) from the culture dish, vigorously pipette the culture medium with 1-mL disposable pipette for 10-20 times [cells remain adhered to dish].
12. Remove the culture medium and immediately replace it with fresh Neurobasal medium with supplements.
13. Change half the Neurobasal medium with supplements every 3 or 4 days. Pipetting to dislodge contaminants is not required.

Electroporation:

1. Coat a 35-mm glass base dish for culture of transfected cells with 2 mL of 0.08 mg/mL Poly-L-Lysine for more than 2 h and then wash the coated dish with 1 mL of PBS twice. Keep the dish aside for later (step 9).
2. Set the NEPA21 parameters.
3. Mix 20 µL of Opti-MEM and 1.0-2.5 µg of plasmids in a 1.5-mL microtube.
4. Collect all the medium from the cell culturing dish and store in a 15-mL centrifuge tube at 37C (referred as conditioned medium).
5. Wash the cells with 2 mL of PBS twice.
6. Add 1.5 mL of 0.25% trypsin-EDTA and observe under microscopy to ensure that the cells are detached.
7. Add 1.5 mL of DMEM to the dish, resuspend by pipetting several times, and transfer to a 15-mL conical tube.
8. Add 1.5 mL of DMEM again to the dish, resuspend by pipetting several times, and transfer to the same 15-mL conical tube.
9. Centrifuge the cell suspension at 1000 rpm for 5 min. Meanwhile, add 1 mL of Neurobasal medium and 1 mL of conditioned medium to the PLL-coated dish (set aside for step 19).
10. Remove the supernatant from the centrifuged sample, add 2 mL of DMEM, and resuspend by pipetting several times.
11. Centrifuge the cell suspension at 1000 rpm for 5 min.
12. Remove the supernatant, add 1 mL of Neurobasal medium, and resuspend by pipetting.
13. Count the cell number using a hemocytometer.
14. Transfer 4.0-6.0 × 105 cells to a 1.5-mL microtube.
15. Centrifuge the microtube at 1000 rpm for 10 min.
16. Remove all the supernatant carefully using a micropipette.
17. Add the mixture of Opti-MEM and plasmids (from step 3) and resuspend by pipetting several times.
18. Transfer the mixture to a NEPA 2 mm cuvette using a micropipette to prevent foaming and then apply voltage.
19. Add a small amount of medium removed from the PLL-coated dish (from step 9) to the cuvette using a NEPA dropper.
20. Transfer the cell suspension in the cuvette to the PLL-coated dish using the NEPA dropper and incubate at 37°C.

Lipofection:

One-Step Transfection:

1. Coat a 35-mm glass base dish for culture of transfected cells with 2 mL of 0.08 mg/mL Poly-L-Lysine for more than 2 h and then wash the coated dish with 1 mL of PBS twice.
2. Collect all the medium from the cell culturing dish and store in a 15-mL conical tube at 4C (referred as conditioned medium).
3. Wash the cultured cell with 1 mL of PBS twice.
4. Add 1 mL of 0.25% trypsin-EDTA and observe under microscopy to ensure that the cells are detached.
5. Add 1 mL of DMEM to the dish, resuspend by pipetting several times, and transfer to a 15-mL conical tube.
6. Add 1 mL of DMEM again to the dish, resuspend by pipetting several times, and transfer to the same 15-mL conical tube.
7. Centrifuge the cell suspension at 1000 rpm for 5 min.
8. Remove the supernatant, add 1 mL of antibiotic-free Neurobasal medium, and resuspend by pipetting.
9. Count the cell number using a hemocytometer.
10. Transfer 0.3-0.9 × 105 cells to a 1.5-mL microtube.
11. Prepare another two 1.5-mL microtubes containing 15 µL of Dilution buffer.
12. Add 25-600 ng of plasmid to one of the microtubes.
13. Add 0.15-1.8 µL of ScreenFect TMA Reagent to the other microtube (ensure the Reagent is vortexed well before use).
14. Combine the plasmid and Reagent solutions, mix immediately by pipetting 10 times, and allow to stand at room temperature for more than 5 min for complex formation.
15. Add the mixture to the microtube containing cells and mix gently by pipetting several times.
16. Transfer the cell suspension in the microtube to a PLL-coated glass base dish and incubate at 37°C.

Two-step Transfection:

1. Prepare two 1.5-mL microtubes containing 15 µL of Dilution buffer.
2. Add 25-600 ng of plasmid to one of the microtubes.
3. Add 0.15-1.8 µL of ScreenFect TMA Reagent to the other microtube (vortex the Reagent well before use).
4. Combine the plasmid and Reagent solutions, mix immediately by pipetting 10 times, and allow to stand at room temperature for 15-20 min for complex formation.
5. Collect all the medium from the cell culturing dish and store in a 15-mL conical tube at 4C (referred as conditioned medium).
6. Wash the cells with 2 mL of PBS once.
7. Add 1 mL of antibiotic-free Neurobasal medium to the mixture of plasmid and Reagent.
8. Add this mixture to the cell culturing dish and gently shake the dish to mix.
9. Add 1 mL of antibiotic-free Neurobasal medium to the cell culturing dish.
10. The next day, replace the entire medium with the conditioned medium warmed to 37°C.

Lentivirus infection:

(Must be performed in a Biosafety Level 2 laboratory)

1. Thaw the virus in cryotubes under running water.
2. Remove all or 1 mL of the medium from the cell culturing dish.
3. Add 0.2, 1, or 2 mL of virus solution to the dish and incubate at 37°C.
4. The next day, replace the entire medium with Neurobasal medium containing antibiotics.

Generation of overlay images of fluorescence images and phase-contrast grid images:

(All steps were performed using ImageJ; see Fig. S4)

1. Capture fluorescence images of the cells (A) along with phase-contrast images of the grid of the glass-bottom dishes (B) at different optical foci but in the same area.
2. Open one phase-contrast image of the grid (B); then, to extract all grid line edges (yellow) within the image, use Wand Tool to click on black grid lines [at the appropriate tolerance setting] (C).
3. Save the extracted grid edges of using ‘ROI manager’ (C).
4. Keep the window of ROI settings open, and then open the cell fluorescence image and generate a binarized fluorescence image of the cells using ‘Threshold’ (D).
5. Extract the extracellular areas of the binarized fluorescence image from (D) using Wand Tool and save these areas in the same ROI setting window using ‘ROI manager’ (E).
6. Select the two ROI saved in steps 3 and 5 in the same ‘ROI manager’ window, and superimpose using the AND command in ‘ROI manager’ (More > AND) to generate grid edge images subtracted with cell area [yellow lines] (F). Arrow in (F) indicates the area of the cell overlapping with the grid that is removed.
7. Select the phase-contrast image with the grid excluding cell areas generated in step 6 and copy (Edit > Copy), and then open the fluorescence image (A) again and paste (Edit > Paste) onto this image, to create a fluorescence image with the grid (G).
8. Save the image generated in step 7 as a new tif file (H).
9. Adjust the positions of fluorescence images captured every 2 h by rotation and cropping according to the grid so that the images at each time are in the same position. The corresponding phase-contrast images of the cells can be created in the same manner of rotation and cropping as the fluorescence images (see Fig. 5F).

## Acknowledgments

We thank Professor Baljit Khakh for the discussions and suggestions on the terminology used in this paper.

## Author contributions

Conceptualization: C.I., K. I., H.N., A.G.T.; Methodology: C.I., K.I., K.K., S.T., A.T., S.N., Y.I., T.T., K.T., A.N., A.G.T.; Formal analysis: C.I., K.I., K.K., T.T.; Investigation: C.I., K.I., K.K., S.T., A.T., S.N.,.; Resources: A.N., K.T., H.N.; Data curation: C.I., K.I., K.K.; Writing - original draft: C.I., A.G.T.; Writing - review & editing: E.S., A.G.T.; Visualization: C.I., K.I., K.K., T.T; Supervision: E.S., K.S, A.G.T.; Project administration: A.G.T.; Funding acquisition: A.G.T.

## Data availability

All relevant data and supplementary information can be found within the article and our previous paper (Tsukuda et al., 2019).

## References

Arizono, M., Idziak, A., Quici, F., & Nägerl, U. V. (2023). Getting sharper: the brain under the spotlight of super-resolution microscopy. Trends in Cell Biology, 33(2), 148–161. 10.1016/j.tcb.2022.06.011

Aumann, G., Friedländer, F., Thümmler, M., Keil, F., Brunkhorst, R., Korf, H. W., & Derouiche, A. (2017). Quantifying Filopodia in Cultured Astrocytes by an Algorithm. Neurochemical Research, 42(6), 1795–1809. 10.1007/s11064-017-2193-0

Badia-Soteras, A., Heistek, T. S., Kater, M. S. J., Mak, A., Negrean, A., van den Oever, M. C., Mansvelder, H. D., Khakh, B. S., Min, R., Smit, A. B., & Verheijen, M. H. G. (2023). Retraction of Astrocyte Leaflets From the Synapse Enhances Fear Memory. Biological Psychiatry, 94(3), 226–238. 10.1016/j.biopsych.2022.10.013

Baldwin, K. T., Murai, K. K., & Khakh, B. S. (2024). Astrocyte morphology. Trends in Cell Biology, 34(7), 547–565. 10.1016/j.tcb.2023.09.006

Ben Haim, L., & Rowitch, D. H. (2016). Functional diversity of astrocytes in neural circuit regulation. Nature Reviews Neuroscience, 18(1), 31–41. 10.1038/nrn.2016.159

Blanco-Suárez, E., Caldwell, A. L. M., & Allen, N. J. (2017). Role of astrocyte–synapse interactions in CNS disorders. Journal of Physiology, 595(6), 1903–1916. 10.1113/JP270988

Butt, E., & Raman, D. (2018). New frontiers for the cytoskeletal protein LASP1. Frontiers in Oncology (Vol. 8, Issue SEP, pp. 1–11). 10.3389/fonc.2018.00391

Derouiche, a, & Frotscher, M. (2001). Peripheral astrocyte processes: monitoring by selective immunostaining for the actin-binding ERM proteins. Glia, 36(3), 330–341. http://www.ncbi.nlm.nih.gov/pubmed/11746770

Derouiche, A., & Geiger, K. D. (2019). Perspectives for ezrin and radixin in astrocytes: Kinases, functions and pathology. International Journal of Molecular Sciences, 20(15). 10.3390/ijms20153776

Endo, F., Kasai, A., Soto, J. S., Yu, X., Qu, Z., Hashimoto, H., Gradinaru, V., Kawaguchi, R., & Khakh, B. S. (2022). Molecular basis of astrocyte diversity and morphology across the CNS in health and disease. Science, 378(6619). 10.1126/science.adc9020

Endo, M., Ohashi, K., & Mizuno, K. (2007). LIM kinase and slingshot are critical for neurite extension. The Journal of Biological Chemistry, 282(18), 13692–13702. 10.1074/jbc.M610873200

Flores, L. R., Keeling, M. C., Zhang, X., Sliogeryte, K., & Gavara, N. (2019). Lifeact-GFP alters F-actin organization, cellular morphology and biophysical behaviour. Scientific Reports, 9(1), 1–13. 10.1038/s41598-019-40092-w

Foo, L. C. (2013). Purification of rat and mouse astrocytes by immunopanning. Cold Spring Harbor Protocols, 8(5), 421–432. 10.1101/pdb.prot074211

Foo, L. C., Allen, N. J., Bushong, E. A., Ventura, P. B., Chung, W. S., Zhou, L., Cahoy, J. D., Daneman, R., Zong, H., Ellisman, M. H., & Barres, B. A. (2011). Development of a method for the purification and culture of rodent astrocytes. Neuron, 71(5), 799–811. 10.1016/j.neuron.2011.07.022

Fujita, Y., Chokki, T., Nishioka, T., Morimoto, K., Nakayama, A., Nakae, H., Ogasawara, M., & Terasaki, A. G. (2021). The emergence of nebulin repeats and evolution of lasp family proteins. Cytoskeleton, 78(9), 419–435. 10.1002/cm.21693

Gallo, G. (2024). The Axonal Actin Filament Cytoskeleton: Structure, Function, and Relevance to Injury and Degeneration. Molecular Neurobiology, 61(8), 5646–5664. 10.1007/s12035-023-03879-7

Geiger, B. (1979). A 130K Protein from Chicken GizzardC: Its Localization at the Termini of Microfilament Bundles in Cultured Chicken Cells. Cell, 18(1), 193–205. 10.1016/0092-8674(79)90368-4

Güler, B. E., Krzysko, J., & Wolfrum, U. (2021). Isolation and culturing of primary mouse astrocytes for the analysis of focal adhesion dynamics. STAR Protocols, 2(4). 10.1016/j.xpro.2021.100954

Güler, B. E., Linnert, J., & Wolfrum, U. (2023). Monitoring paxillin in astrocytes reveals the significance of the adhesion G protein coupled receptor VLGR1/ADGRV1 for focal adhesion assembly. Basic and Clinical Pharmacology and Toxicology, 133(4), 301–312. 10.1111/bcpt.13860

Haseleu, J., Anlauf, E., Blaess, S., Endl, E., & Derouiche, A. (2013). Studying subcellular detail in fixed astrocytes: dissociation of morphologically intact glial cells (DIMIGs). Frontiers in Cellular Neuroscience, 7(May), 1–10. 10.3389/fncel.2013.00054

Holt, L. M., Hernandez, R. D., Pacheco, N. L., Torres Ceja, B., Hossain, M., & Olsen, M. L. (2019). Astrocyte morphogenesis is dependent on BDNF signaling via astrocytic TrkB.T1. ELife, 8, 1–27. 10.7554/eLife.44667

John, G. R. (2004). Interleukin-1 Induces a Reactive Astroglial Phenotype via Deactivation of the Rho GTPase-Rock Axis. Journal of Neuroscience, 24(11), 2837–2845. 10.1523/JNEUROSCI.4789-03.2004

Kálmán, M., Székely, A. D., & Csillag, A. (1998). Distribution of glial fibrillary acidic protein and vimentin-immunopositive elements in the developing chicken brain from hatch to adulthood. Anatomy and Embryology, 198(3), 213–235. 10.1007/s004290050179

Kanchanawong, P., & Calderwood, D. A. (2023). Organization, dynamics and mechanoregulation of integrin-mediated cell-ECM adhesions. Nature Reviews Molecular Cell Biology volume, 24(February). 10.1038/s41580-022-00531-5

Lange, S. C., Bak, L. K., Waagepetersen, H. S., Schousboe, A., & Norenberg, M. D. (2012). Primary cultures of astrocytes: Their value in understanding astrocytes in health and disease. Neurochemical Research, 37(11), 2569–2588. 10.1007/s11064-012-0868-0

Lavialle, M., Aumann, G., Anlauf, E., Pröls, F., Arpin, M., & Derouiche, A. (2011). Structural plasticity of perisynaptic astrocyte processes involves ezrin and metabotropic glutamate receptors. Proceedings of the National Academy of Sciences of the United States of America, 108(31), 12915–12919. 10.1073/pnas.1100957108

Lawrence, J. M., Schardien, K., Wigdahl, B., & Nonnemacher, M. R. (2023). Roles of neuropathology-associated reactive astrocytes: a systematic review. Acta Neuropathologica Communications, 11(1), 1–28. 10.1186/s40478-023-01526-9

Liu, J. S. (2011). Molecular genetics of neuronal migration disorders. Current Neurology and Neuroscience Reports, 11(2), 171–178. 10.1007/s11910-010-0176-5

Lowery, L. A., & Van Vactor, D. (2009). The trip of the tip: understanding the growth cone machinery. Nature Reviews. Molecular Cell Biology, 10(5), 332–343. 10.1038/nrm2679

Molotkov, D., Zobova, S., Arcas, J. M., & Khiroug, L. (2013). Calcium-induced outgrowth of astrocytic peripheral processes requires actin binding by Profilin-1. Cell Calcium, 53(5– 6), 338–348. 10.1016/j.ceca.2013.03.001

Moye, S. L., Diaz-Castro, B., Gangwani, M. R., & Khakh, B. S. (2019). Visualizing astrocyte morphology using lucifer yellow iontophoresis. Journal of Visualized Experiments, (151), 1–10. 10.3791/60225

Nakagawa, H., Suzuki, H., Machida, S., Suzuki, J., Ohashi, K., Jin, M., Miyamoto, S., & Terasaki, A. G. (2009). Contribution of the LIM domainand nebulin-repeatsto the interaction of Lasp-2 with actin filaments and focal adhesions. PLoS ONE, 4(10), 8–13. 10.1371/journal.pone.0007530

Oberheim, N. A., Takano, T., Han, X., He, W., Lin, J. H. C., Wang, F., Xu, Q., Wyatt, J. D., Pilcher, W., Ojemann, J. G., Ransom, B. R., & Goldman, S. A. (2009). Uniquely Hominid Features of Adult Human Astrocytes. Journal of Neuroscience, 29(10), 3276–3287. 10.1523/JNEUROSCI.4707-08.2009

Octeau, J. C., Chai, H., Jiang, R., Bonanno, S. L., Martin, K. C., & Khakh, B. S. (2018). An Optical Neuron-Astrocyte Proximity Assay at Synaptic Distance Scales. Neuron, 98(1), 49–66.e9. 10.1016/j.neuron.2018.03.003

Ohta, K., Higashi, R., Sawaguchi, A., & Nakamura, K. ichiro. (2012). Helical arrangement of filaments in microvillar actin bundles. Journal of Structural Biology, 177(2), 513–519. 10.1016/j.jsb.2011.10.012

Panda, S. K., & McGrew, M. J. (2022). Genome editing of avian species: implications for animal use and welfare. Laboratory Animals, 56(1), 50–59. 10.1177/0023677221998400

Pettmann, B., Louis, J. & Sensenbrenner, M. (1979) Morphological and biochemical maturation of neurones cultured in the absence of glial cells. Nature 281, 378–380. 10.1038/281378a0

Riedl, J., Crevenna, A. H., Kessenbrock, K., Yu, J. H., Neukirchen, D., Bista, M., Bradke, F., Jenne, D., Holak, T. A., Werb, Z., Sixt, M., & Wedlich-Soldner, R. (2008). Lifeact: a versatile marker to visualize F-actin. Nature Methods, 5(7), 605–607. 10.1038/nmeth.1220

Saint-Martin, M., & Goda, Y. (2022). Astrocyte–synapse interactions and cell adhesion molecules. FEBS Journal, 290(14), 1–15. 10.1111/febs.16540

Saura, J. (2007). Microglial cells in astroglial cultures: A cautionary note. Journal of Neuroinflammation, 4, 1–11. 10.1186/1742-2094-4-26

Schacke, S., Kirkpatrick, J., Stocksdale, A., Bauer, R., Hagel, C., Riecken, L. B., & Morrison, H. (2022). Ezrin deficiency triggers glial fibrillary acidic protein upregulation and a distinct reactive astrocyte phenotype. Glia, 70(12), 2309–2329. 10.1002/glia.24253

Schiweck, J., Eickholt, B. J., & Murk, K. (2018). Important shapeshifter: Mechanisms allowing astrocytes to respond to the changing nervous system during development, injury and disease. Frontiers in Cellular Neuroscience, 12(August), 1–17. 10.3389/fncel.2018.00261

Semyanov, A., & Verkhratsky, A. (2021). Astrocytic processes: from tripartite synapses to the active milieu. Trends in Neurosciences, 44(10), 781–792. 10.1016/j.tins.2021.07.006

Shigetomi, E., Patel, S., & Khakh, B. S. (2016). Probing the Complexities of Astrocyte Calcium Signaling. Trends in Cell Biology, 26(4), 300–312. 10.1016/j.tcb.2016.01.003

Smith, J., Alfieri, J. M., Anthony, N., Arensburger, P., Athrey, G. N., Balacco, J., Balic, A., Bardou, P., Barela, P., Bigot, Y., Blackmon, H., Borodin, P. M., Carroll, R., Casono, M. C., Charles, M., Cheng, H., Chiodi, M., Cigan, L., Coghill, L. M., … Zhou, H. (2023). Fourth Report on Chicken Genes and Chromosomes 2022. Cytogenetic and Genome Research, 162(8–9), 405–528. 10.1159/000529376

Smith, M. A., Hoffman, L. M., & Beckerle, M. C. (2014). LIM proteins in actin cytoskeleton mechanoresponse. Trends in Cell Biology, 24(10), 575–583. 10.1016/j.tcb.2014.04.009

Soto, J. S., Jami-Alahmadi, Y., Chacon, J., Moye, S. L., Diaz-Castro, B., Wohlschlegel, J. A., & Khakh, B. S. (2023). Astrocyte–neuron subproteomes and obsessive–compulsive disorder mechanisms. Nature (Vol. 616, Issue 7958). Springer US. 10.1038/s41586-023-05927-7

Soto, J. S., Neupane, C., Kaur, M., Pandey, V., Wohlschlegel, J. A., & Khakh, B. S. (2024). Astrocyte Gi-GPCR signaling corrects compulsive-like grooming and anxiety-related behaviors in Sapap3 knockout mice. Neuron, 112. 10.1016/j.neuron.2024.07.019

Svitkina, T. (2018). The actin cytoskeleton and actin-based motility. Cold Spring Harbor Perspectives in Biology, 10(1). 10.1101/cshperspect.a018267

Terasaki, A. G., Suzuki, H., Nishioka, T., Matsuzawa, E., Katsuki, M., Nakagawa, H., Miyamoto, S., & Ohashi, K. (2004). A novel LIM and SH3 protein (lasp-2) highly expressing in chicken brain. Biochemical and Biophysical Research Communications, 313(1), 48–54. 10.1016/j.bbrc.2003.11.085

Thomason, E. J., Escalante, M., Osterhout, D. J., & Fuss, B. (2020). The oligodendrocyte growth cone and its actin cytoskeleton: A fundamental element for progenitor cell migration and CNS myelination. Glia, 68(7), 1329–1346. 10.1002/glia.23735

Torres-Ceja, B., & Olsen, M. L. (2022). A closer look at astrocyte morphology: Development, heterogeneity, and plasticity at astrocyte leaflets. Current Opinion in Neurobiology, 74, 102550. 10.1016/j.conb.2022.102550

Tsukuda, S., Tamura, A., Matsumoto, K., Fujita, Y., Nakata, S., Kaneko, T., Nakayama, A., Nakagawa, H., & Terasaki, A. G. (2019). In vitro differentiation of chicken astrocytes: Growth, morphology, and protein expression of astrocytes in primary cultures. Zoological Science, 36(6). 10.2108/zs180102

Wilson, D. M., Cookson, M. R., Van Den Bosch, L., Zetterberg, H., Holtzman, D. M., & Dewachter, I. (2023). Hallmarks of neurodegenerative diseases. Cell, 186(4), 693–714. 10.1016/j.cell.2022.12.032

Yu, X., & Khakh, B. S. (2022). SnapShot: Astrocyte interactions. Cell, 185(1), 220–220.e1. 10.1016/j.cell.2021.09.029

Zieseniss, A., Terasaki, A. G., & Gregorio, C. C. (2008). Lasp-2 expression, localization, and ligand interactions: A new Z-disc scaffolding protein. Cell Motility and the Cytoskeleton, 65(1), 59–72. 10.1002/cm.20244

