## Supplementary material for "Improved transfection methods of primary cultured astrocytes for observation of cytoskeletal structures": Sup Figs

Fig.S1

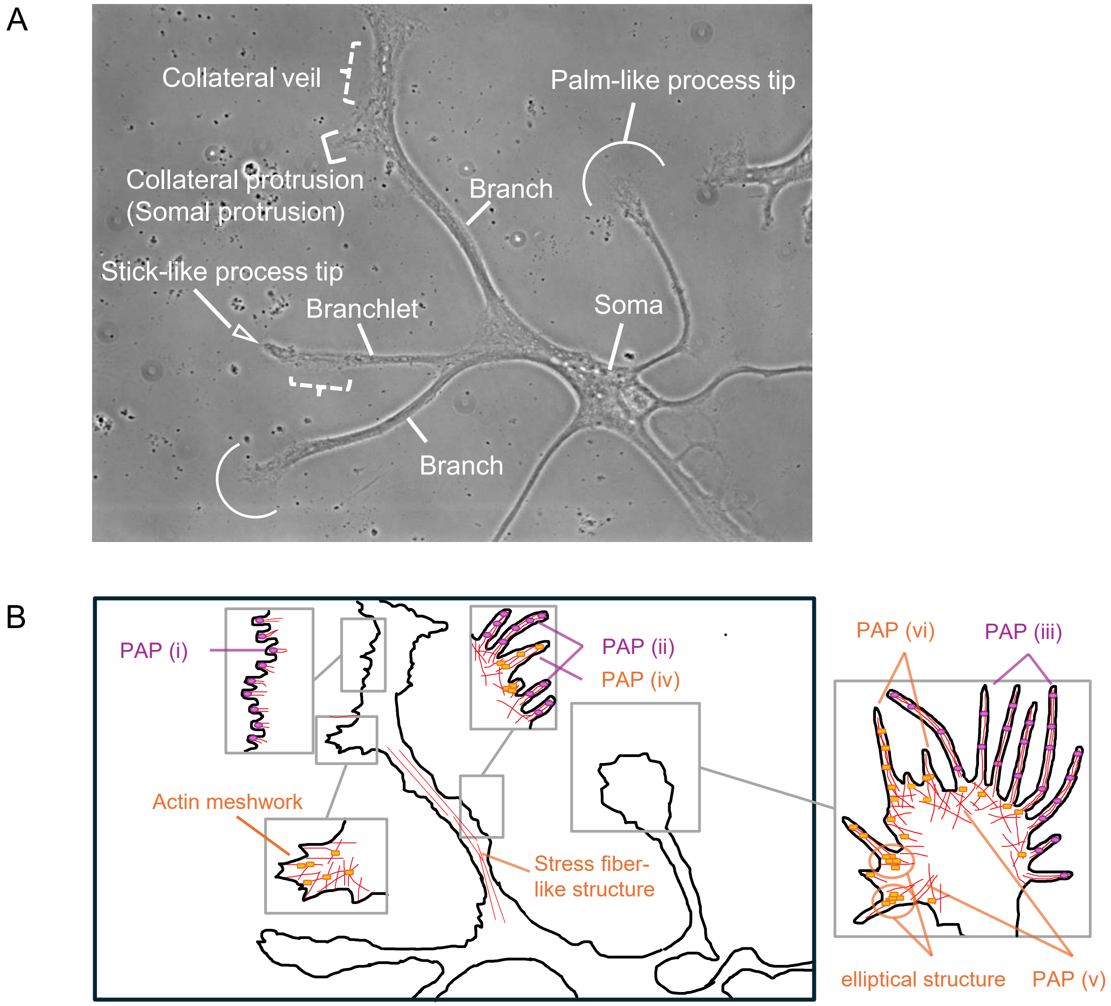


C

| PAP-like structures in cell periphery | ezrin | lasp-2 |
|---|---|---|
| (i) microvilli at the edges of collateral veils | + | - |
| (ii) microvilli directly extending from stem process | + | - |
| (iii) microvilli at process tips | + | - |
| (iv) filopodia directly extending from stem process | - | + |
| (v) lamellipodia at process tips | - | + |
| (vi) filopodia at process tips | - | + |
| **Other actin-containing structures in cell periphery** | **ezrin** | **lasp-2** |
| actin meshwork in somal protrusions | - | + |
| actin meshwork in collateral protrusions | - | + |
| **Actin-containing structures in cell inner** | **ezrin** | **lasp-2** |
| elliptical structures at process tips | - | + |
| stress fiber-like structures | - | - |

Fig.S1

**Fig. S1. Terminology of structures in cultured astrocytes and summary of ezrin and lasp-2 localization in actin-containing structures**

(A) Terminology of structures of cultured astrocytes in the current study. “Soma”: cell body; “Branch”: major stem process more than 10 μm in length emanating from the astrocyte soma; “Branchlet”: secondary/tertiary stem process from branch; “Stem process”: covers all processes including branches and branchlets; “Somal protrusion” and “Collateral protrusion” (square bracket): a 5-10 μm protoplasmic protrusion (not like filopodia and lamellipodia) directly extended from soma, or, at the side of a stem process, expected to grow into a branch or a branchlet; “Collateral veil” (dashed curly bracket): thin membrane ruffles that are difficult to identify in phase-contrast images, but clearly stained with anti-ezrin antibody (see Fig. 3A); "Process tip": tip of a stem process indicated by either arrowhead (stick-like structure with process width < 2 times the branch/branchlet width) or arc (larger palm-like structure with process width > 2 times the branch/branchlet width). In many cases, the distinction between branch and branchlet is rather arbitrary (branchlet here could also be considered a branch and vice versa). Scale bar = 20 μm.

(B) Schematic illustration of observed actin-containing structures corresponding to phase-contrast image of (A). In phase-contrast images, it is difficult to observe finer structures, but enlarged images of immunostaining or transfected fluorescent vectors could identify fine structures of actin cytoskeleton (see enlarged boxes). "PAP" with Roman numerals indicate the six types of PAP-like structures classified in the current study. Red lines: actin filaments; purple circles: ezrin; orange squares: lasp-2.

(C) Tabular summary of ezrin and lasp-2 localization in actin-containing structures. +, localized; -, not localized

Fig.S2

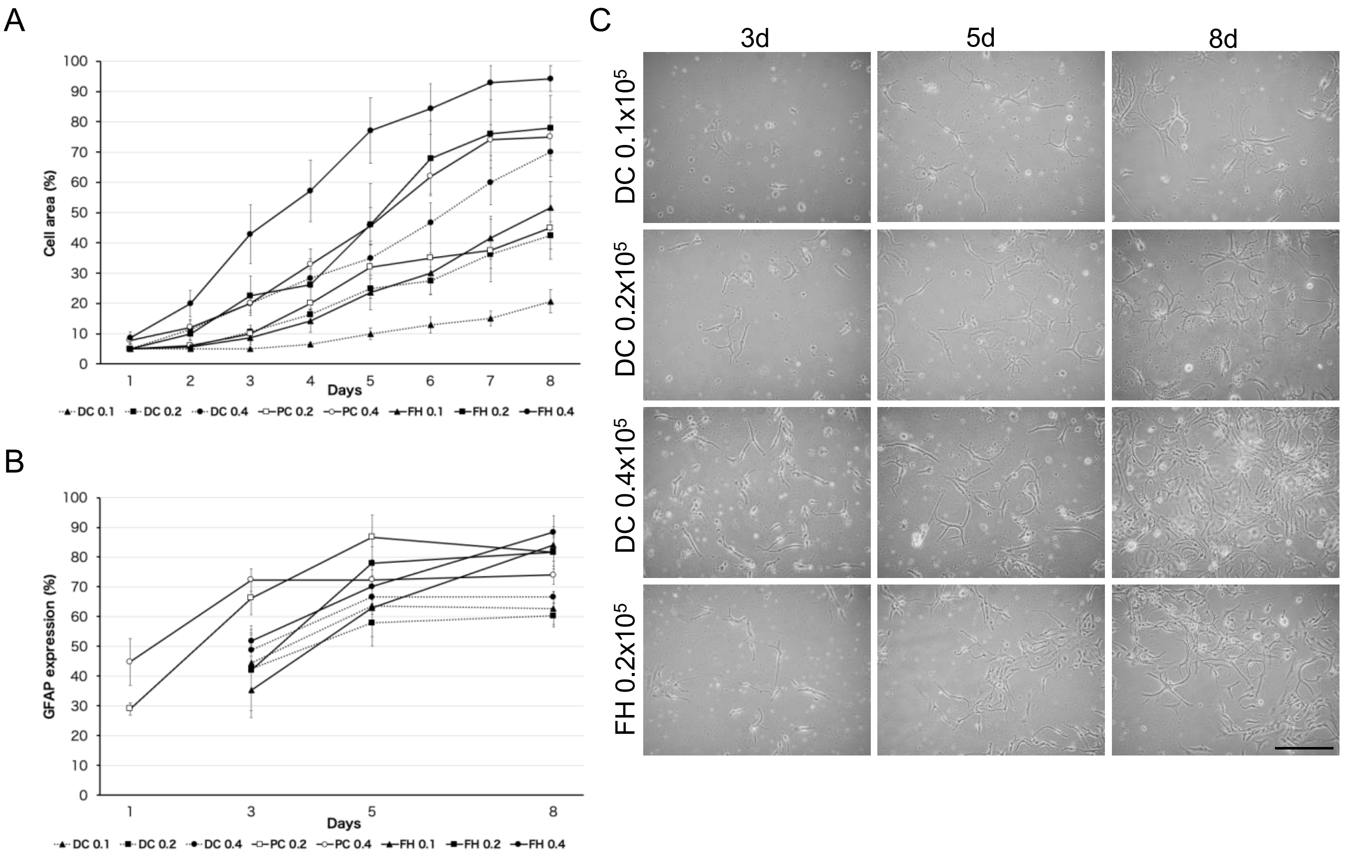


**Fig. S2. Proliferation and differentiation of directly cryopreserved astrocytes**
(A) Cell coverage area of freshly harvested (FH) astrocytes, pre-cultured cryopreserved (PC) astrocytes, and directly cryopreserved (DC) astrocytes at different initial cell densities: 0.1, 0.2, and $0.4 \times 10^5$ cells/$cm^2$. The data of FH and PC astrocytes of (A) and (B) are cited from (Tsukuda et al., 2019), and the graph displays the means of independent experiments (N = 5-9) ± SEM. (B) Glial fibrillary acidic protein (GFAP) expression rate of FH, PC, and DC astrocytes for the different initial cell densities. Note that all data from FH and PC astrocytes for comparison with DC astrocytes are from Tsukuda et al. (2019). (C) Representative images of DC astrocytes at different initial cell densities after 3, 5, and 8 days. FH astrocytes at $0.2 \times 10^5$ cells/$cm^2$ initial cell density is used as control. Scale bar = 200 µm.

Fig.S3

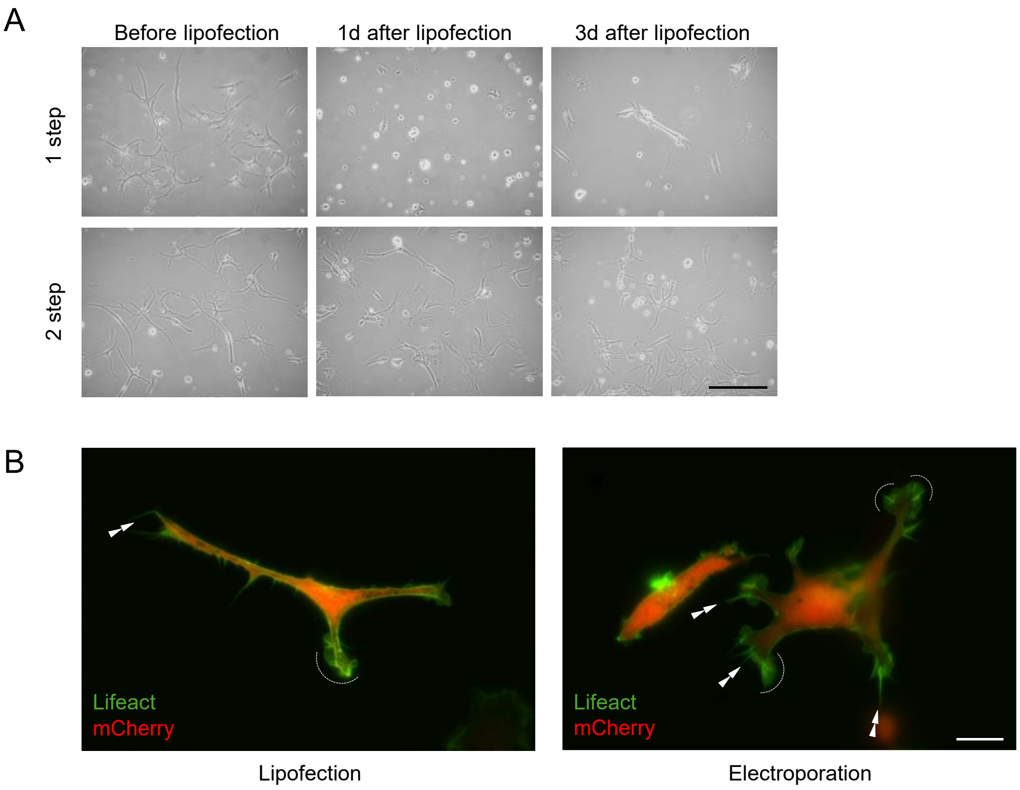


**Fig. S3. Morphology of lipofected astrocytes and localization of EGFP-lifeact in lipofection and electroporation**

(A) Lower magnification view of astrocytes before and after (1d and 3d) transfection with two methods (1 step or 2 step). Cells are cultured at a density of $0.2 \times 10^5$ cells/cm$^2$ for 7 days before lipofection. Scale bar = 200 µm. (B) Representative images of pEGFP-lifeact and pmCherry in astrocytes after 2d of lipofection and electroporation. Double arrowheads, dashed arcs indicate filopodia and lamellipodia at process tips, respectively. Scale bar = 20 µm.

Fig.S4

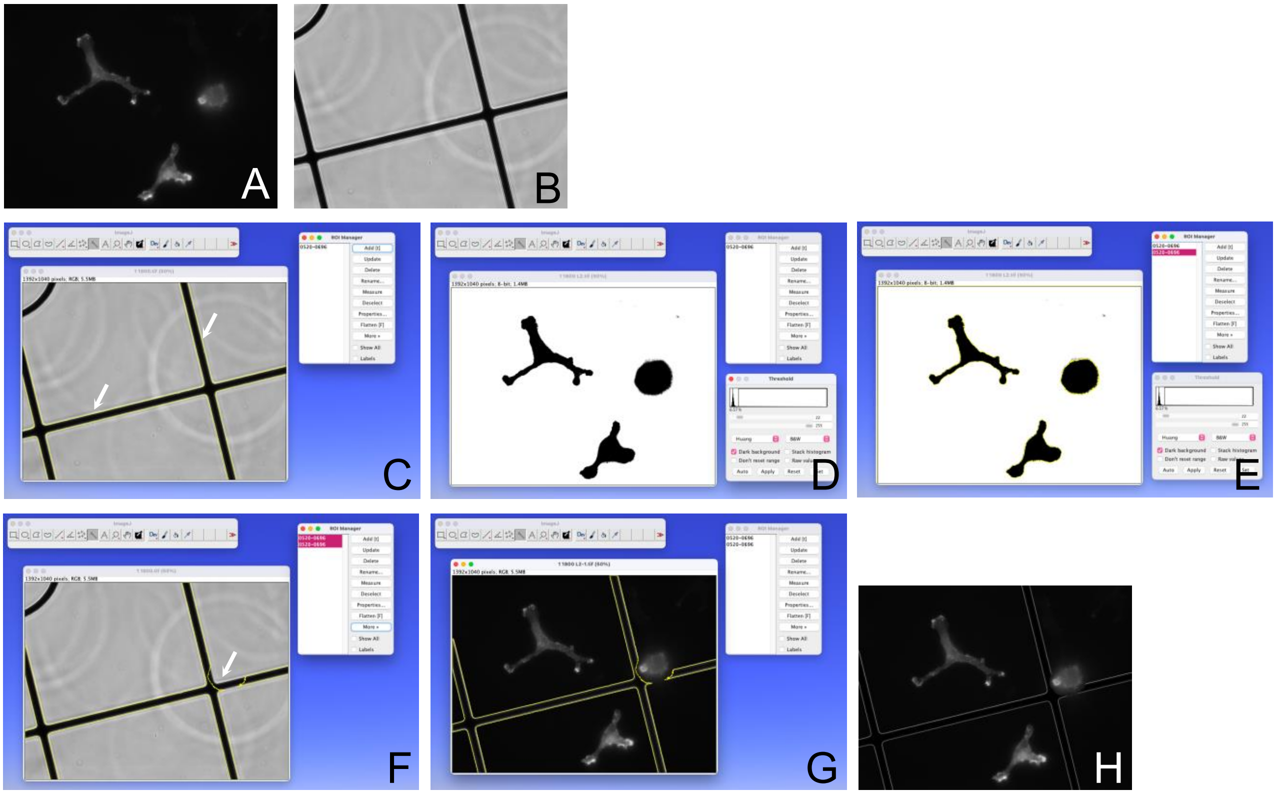


**Fig. S4. Steps of overlaying images of fluorescence images and phase-contrast images**(A) Fluorescence image of cells. (B) Phase-contrast image of grid. (C) Extraction of all grid edges within the image. Arrows indicated extracted grid edges with pale yellow. (D) Binarized fluorescence image. (E) Extraction of extracellular areas of the binarized fluorescence image using Wand Tool. (F) Subtraction of grid edge image with cell area. An arrow indicates a subtracted grid edge. (G) Overlaying of grid edge image on fluorescence image. (H) Completed fluorescence image with the grid.

Sup. movie 1

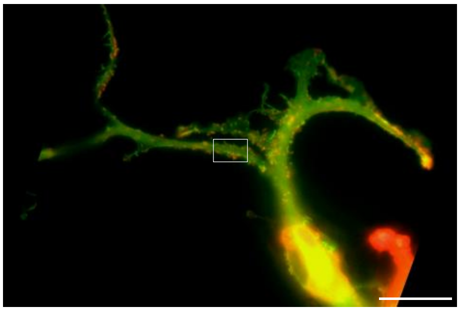

**Supplemental Movie 1**
Movie compiling fluorescent images of astrocytes in Fig. 4D every 20 sec for 2 min. Scale bar = 20 μm.

Sup. movie 2

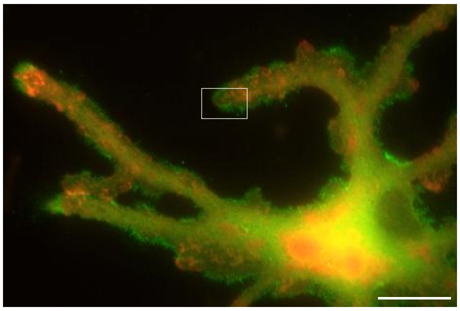

**Supplemental Movie 2**
Movie compiling fluorescent images of astrocytes in Fig. 5F every 20 sec for 3 min. Scale bar = 20 μm.
